# Discovery of S-217622, a Non-Covalent Oral SARS-CoV-2 3CL Protease Inhibitor Clinical Candidate for Treating COVID-19

**DOI:** 10.1101/2022.01.26.477782

**Authors:** Yuto Unoh, Shota Uehara, Kenji Nakahara, Haruaki Nobori, Yukiko Yamatsu, Shiho Yamamoto, Yuki Maruyama, Yoshiyuki Taoda, Koji Kasamatsu, Takahiro Suto, Kensuke Kouki, Atsufumi Nakahashi, Sho Kawashima, Takao Sanaki, Shinsuke Toba, Kentaro Uemura, Tohru Mizutare, Shigeru Ando, Michihito Sasaki, Yasuko Orba, Hirofumi Sawa, Akihiko Sato, Takafumi Sato, Teruhisa Kato, Yuki Tachibana

## Abstract

The coronavirus disease 2019 (COVID-19) pandemic, caused by severe acute respiratory syndrome coronavirus 2 (SARS-CoV-2), has resulted in millions of deaths and threatens public health and safety. Despite the rapid global spread of COVID-19 vaccines, effective oral antiviral drugs are urgently needed. Here, we describe the discovery of S-217622, the first oral non-covalent, non-peptidic SARS-CoV-2 3CL protease inhibitor clinical candidate. S-217622 was discovered *via* virtual screening followed by biological screening of an in-house compound library, and optimization of the hit compound using a structure-based drug-design strategy. S-217622 exhibited antiviral activity *in vitro* against current outbreaking SARS-CoV-2 variants and showed favorable pharmacokinetic profiles *in vivo* for once-daily oral dosing. Furthermore, S-217622 dose-dependently inhibited intrapulmonary replication of SARS-CoV-2 in mice, indicating that this novel non-covalent inhibitor could be a potential oral agent for treating COVID-19.

## Introduction

The global coronavirus disease 2019 (COVID-19) pandemic, caused by severe acute respiratory syndrome coronavirus 2 (SARS-CoV-2), continues to spread worldwide; more than 315 million people have been infected, and 5.5 million have died as of January 2022.^1^ Because therapeutic options remain limited, oral COVID-19 therapeutics are urgently needed, especially for non-hospitalized patients, to prevent hospitalization and death.^2^

SARS-CoV-2 is highly pathogenic to older adults and persons with high risk factors and can develop into severe, life-threatening acute respiratory distress syndrome. SARS-CoV-2 is an enveloped positive-sense single-stranded RNA virus that is a member of the genus *Betacoronavirus*.^3^ SARS-CoV-2 enters host cells by binding its spike glycoprotein to angiotensin-converting enzyme 2 (ACE2) and releases its viral RNA genome into the cytoplasm after uncoating. After entry, the viral RNA genome subjects the cell to translation of two large polyproteins, pp1a and pp1ab, which are processed into individual non-structural proteins. Nsp5, also known as 3C-like protease (3CL^pro^) or the main protease, is a cysteine protease responsible for cleaving 11 distinct sites of the polyproteins to transform into mature functional proteins. 3CL^pro^ plays a critical role in viral replication, and its inhibition prevents formation of replication-essential enzymes, such as RNA-dependent RNA polymerase, thus inhibiting viral replication.^4^ Viral proteases are well-validated drug targets for treating human immunodeficiency virus and hepatitis C virus and have been used in various approved oral drugs.^5^ Additionally, the antiviral efficacy of the 3CL^pro^ inhibitor would likely be unaffected by and not induce mutations of the spike protein, which often occur in SARS-CoV-2 variants, because the 3CL^pro^ and spike protein are distinct proteins encoded in different regions of the viral genome. Thus, 3CL^pro^ is an attractive target for small-molecule oral therapeutics for treating COVID-19. Recent reports have revealed that peptide-like 3CL^pro^ inhibitors with reactive “warheads” show potent antiviral activities *in vitro*, and some of these drugs reduced viral loads *in vivo* in SARS-CoV-2-infected human ACE2 transgenic mouse models.^6, 7^ Recently, Pfizer reported good results from a clinical study of the peptidic, covalent oral 3CL^pro^ inhibitor, PF-07321332, which is dosed with ritonavir as a pharmacokinetic (PK) booster.^8^ However, challenges remain for improving the target selectivity and PK profiles of peptide-like covalent inhibitors owing to the intrinsic nature of the reactivity, low membrane permeability and low metabolic stability.^9–11^ Hence, non-peptidic, non-covalent small-molecule inhibitors have attracted much attention; however, their potency and PK profiles must be further optimized.^12–16^

Here, we describe the discovery of S-217622, the first non-peptidic, non-covalent SARS-CoV-2 3CL^pro^ inhibitor clinical candidate for treating COVID-19, and its preclinical characterization. S-217622 displayed antiviral activity *in vitro* towards a range of SARS-CoV-2 variants and coronavirus families, favorable drug metabolism and pharmacokinetic (DMPK) profiles for the oral agents, and dose-dependent antiviral efficacy *in vivo*, indicating its potential for once-daily oral treatment of COVID-19.

## Results and Discussions

To rapidly obtain the non-covalent SARS-CoV-2 3CL^pro^ inhibitor clinical candidate to combat with the pandemic, we used a structure-based drug design (SBDD) strategy, starting with docking-based virtual screening followed by biological screening using an in-house compound library (Figure 1). First, we investigated pharmacophores in the binding site of 3CL^pro^ based on the interactions of known inhibitors because applying the pharmacophore filter to the docking screening helps enrich the virtual-screening hit rate.^17^ 3CL^pro^ is a cysteine protease with a Cys145-His41 catalytic dyad in its active site, which strongly recognizes P1 Gln and P2 Leu/Met/Phe/Val as its substrates.^6^ These substrate-like substructures are shared with the potent peptide-like inhibitors, GC-376^7^ and N3^18^, a Gln-mimic lactam moiety in the S1 pocket and Leu-mimic hydrophobic moiety in the S2 pocket (Figure 2a). Non-covalent small-molecule inhibitors, such as ML188,^12^ ML300,^13, 14^ and the 3-aminopyridine-like compound of the Postera COVID moonshot project,^15, 16^ exhibit similar pharmacophores, which form a hydrogen bond with the side-chain NH donor of His163 in the S1 pocket and have fitted lipophilic moieties in the S2 pocket (Figure 2b, c). Additionally, the hydrogen bond with the Glu166 main-chain NH recognizes the P2 main-chain carbonyl of the substrate and is conserved in the known inhibitors. Given these interactions, we hypothesized that these three pharmacophores, i.e., the acceptor site with the side-chain NH donor of His163 in the S1 pocket, the lipophilic site in the S2 pocket, and the acceptor site with the Glu166 main-chain NH, play critical roles in small-molecule binding (Figure 2d).

**Figure 1.**
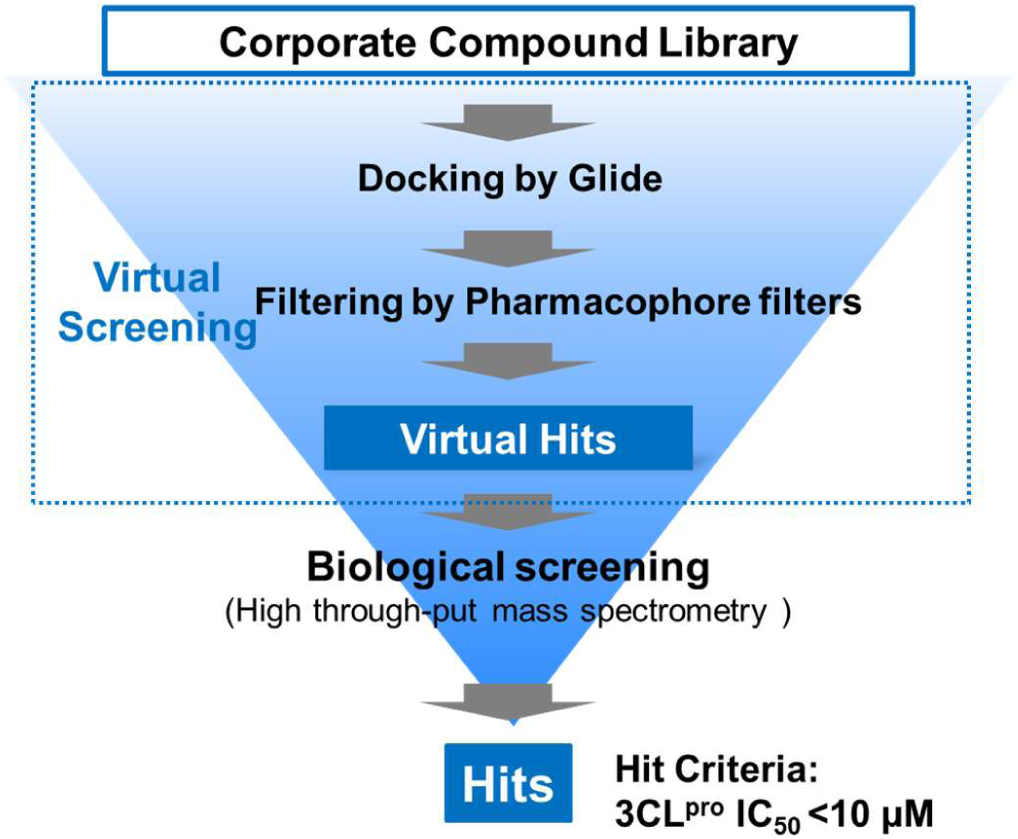
Schematic flow of the screening campaign.

**Figure 2.**
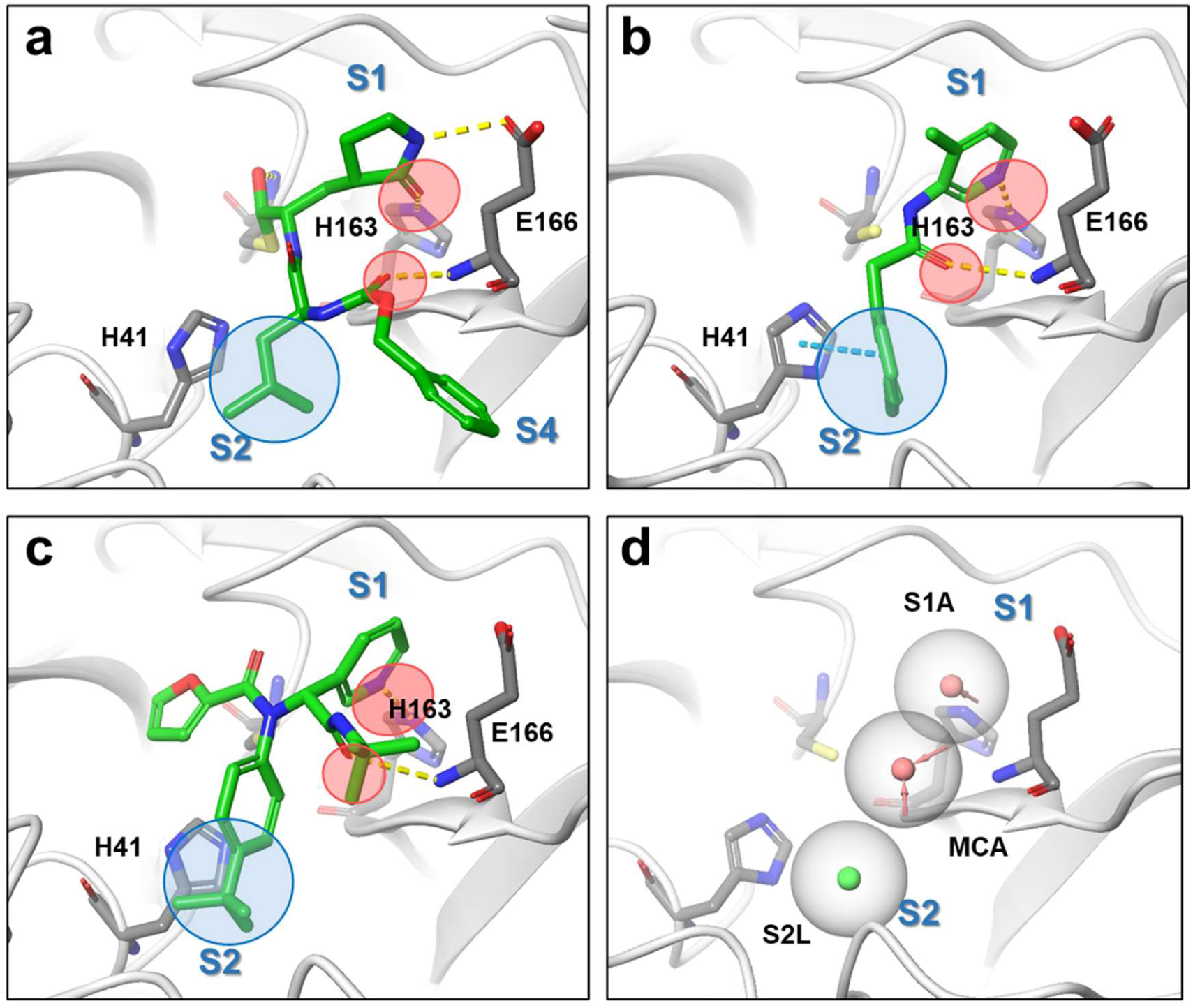
Binding modes of 3CL^pro^ inhibitors, their pharmacophores, and defined pharmacophore filters for virtual screening. (a) Crystal structures of GC376 (PDB: 6WTT), (b) 3-aminopyridine-like compound of the Postera COVID moonshot project (PDB: 5RH2) and (c) ML188 (PDB: 7L0D). The common H-bond acceptors are circled in red; the common hydrophobic pharmacophores are circled in blue. (d) Common pharmacophores shared with inhibitors A–C. Red and green spheres represent H-bond acceptors and lipophilic features, respectively.

We performed docking-based virtual screening using the crystal structures of the 3CL^pro^ and ML188-like non-covalent small molecules (Protein Data Bank [PDB] code: 6w63).^19^ Compounds from the in-house library were docked, then the pharmacophore filter described above was applied to each docking pose, and the 300 top-scoring compounds were evaluated *via* enzymatic assays using mass spectrometry to avoid the false positives that frequently occur in fluorescence-based assays, giving some hit compounds with IC_50_ < 10 μM.

Optimization of the PK profile is a common challenge in drug discovery and usually takes time to overcome. Therefore, if possible, potency optimization of a hit compound with favorable PK profiles is likely the most straightforward way to meet the urgent need for an oral 3CL^pro^ inhibitor. Further profiling of hit compounds revealed that one of the hit compounds, **1**, could be a potential lead for this project because it displayed potent enzymatic inhibitory activity and favorable PK profiles with oral bioavailability (Figure 3). An enzymatic inhibition assay revealed that the IC_50_ value of **1** was 8.6 µM, and the *in vitro* metabolic stabilities of **1,** measured after 30 min of incubation in human and rat microsomes, were 97% and 71%, respectively. An *in vivo* PK study in rats demonstrated that **1** had a favorable profile for the oral agent, oral bioavailability (*F*) of 111% and a low clearance of 7.3 mL/min/mg.

**Figure 3.**
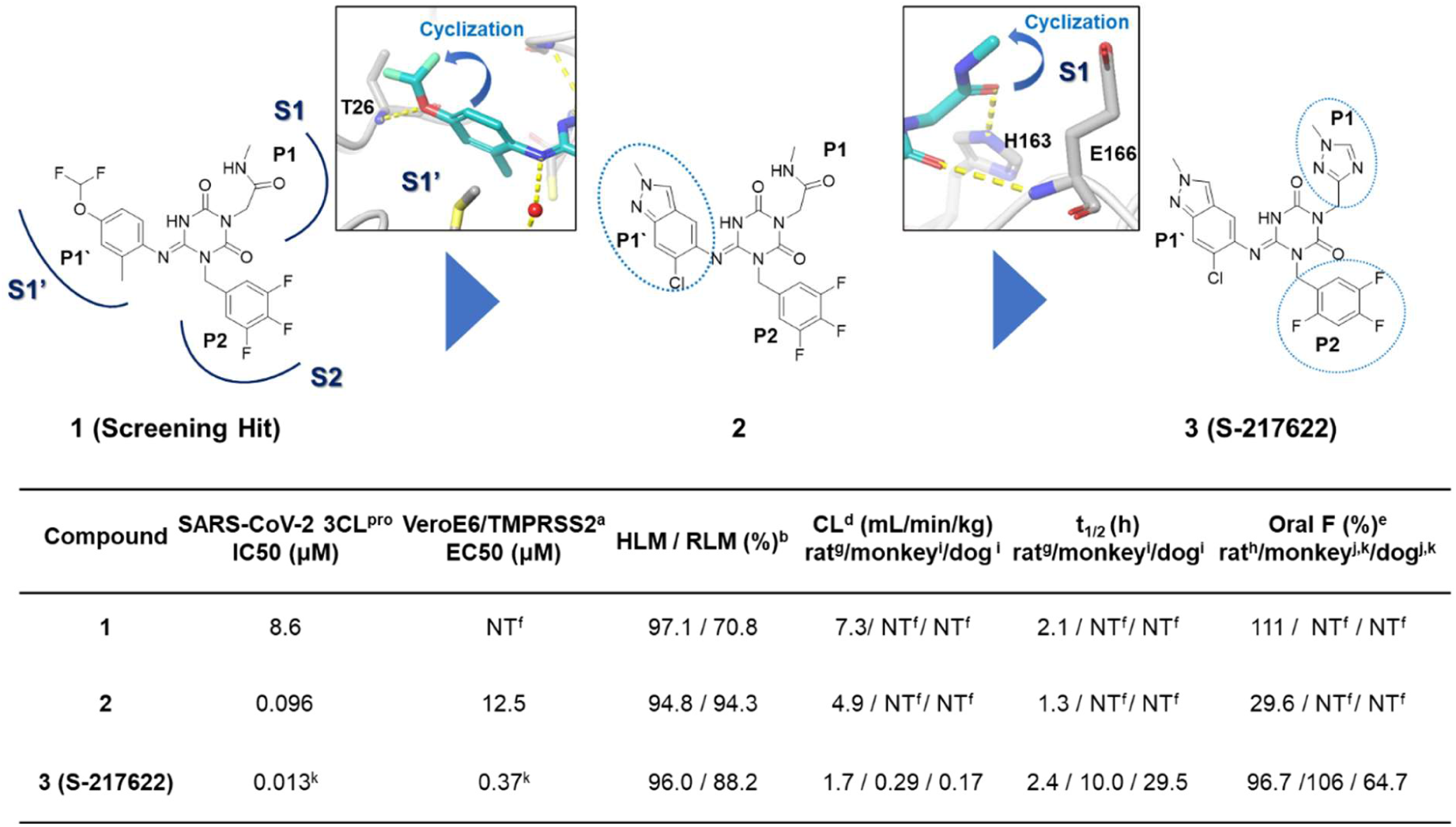
Structure-based optimization of hit compound **1** and profiles of the compounds. ^a^Cytopathic effect inhibition assay with Vero E6 cells expressing human transmembrane protease serine 2 (VeroE6/TMPRSS2). ^b^% remaining in human liver microsomes (HLMs) after 30 min. ^c^% remaining in rat liver microsomes (RLMs) after 30 min. ^d^Total clearance, ^e^Oral bioavailability. ^f^Not tested. ^g^Intravenously administered 0.5 µmol/mL/kg (n = 2), non-fasted; ^h^Orally administered 1 µmol/5 mL/kg (n = 2), non-fasted; ^i^Intravenously administered 0.1 mg/0.2 mL/kg (n = 2), non-fasted; ^j^Orally administered 3 mg/2 mL/kg (n = 3), non-fasted. ^k^Evaluated as S-217622 fumaric acid co-crystal form.

We resolved the X-ray complex structure of **1** with the protease (Figure 4a). As expected, the binding mode of **1** in the X-ray structure was similar to that obtained in the docking (Figure 4b). The S1 and S2 pockets were filled with the methyl-amide and 3,4,5-trifluorobenzene moieties, respectively. The 4-difluoromethoxy-2-methylbenzene subunit was placed in the S1’ pocket. The 2-carbonyl oxygen of the center triazine moiety formed a hydrogen bond with the main-chain NH of Glu166. The other side of the 4-carbonyl oxygen was bound in the oxyanion hole of the protease, which formed two hydrogen bonds with the main-chain NHs of Gly143 and Cys145. The methyl-amide moiety was placed in the S1 pocket, of which, the carbonyl oxygen interacted with the side-chain NH of His163. The protease exhibited an interesting conformational change in the S2 pocket; the side chain of the catalytic His41 was rotated and formed a face-to-face π interaction with the 3,4,5-trifluorobenzene moiety of **1**, whereas the docking pose predicted an edge-to-face π interaction (Figure 4c,d). Along with the side-chain flip of His41, the 4-difluoromethoxy-2-methylbenzene fragment was placed in a slightly different site compared with that of the docking pose, in which the ether oxygen of the P1’ ligand formed a hydrogen bond with the main-chain NH of Thr26. An imine linker formed a water-mediated hydrogen bond with the His41 side chain, indicating its contribution to the affinity.

**Figure 4.**
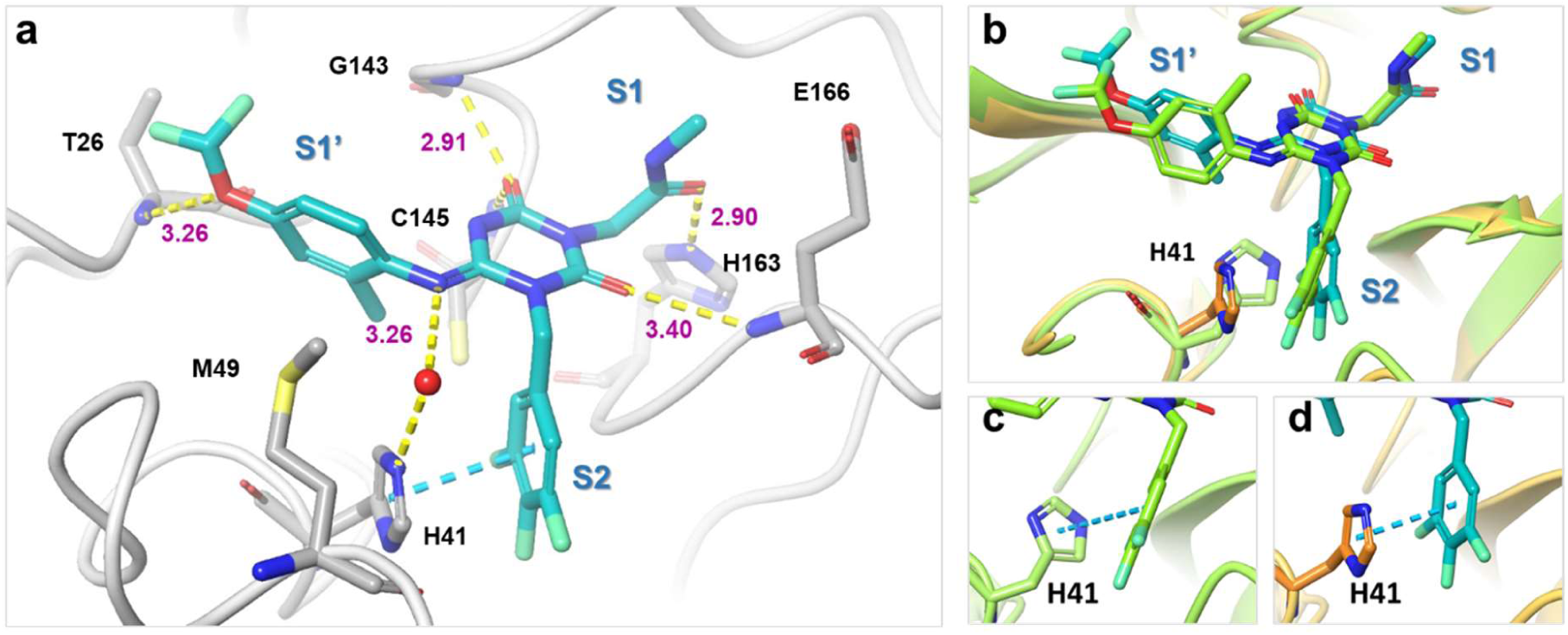
X-ray co-structure of hit compound **1** and 3CL^pro^. (a) Close-up view of **1** (cyan) in the binding pocket. Water molecules are shown as red spheres. Hydrogen bonds are indicated as yellow dashed lines; π-π stacking is indicated as a cyan dashed line. (b-d) Comparison of a docking pose and X-ray crystal structure of **1**. Near the S2 pocket, the sidechain of His41 was rotated to form a face-to-face π interaction with the 3,4,5-trifluorobenzene moiety of **1**. Docked structure is in lime green, and the X-ray structure is in cyan (**1**) and orange (protein residues).

Keeping the hydrogen bonds confirmed by the X-ray complex structure, straightforward multiparameter optimization was achieved starting from hit compound **1** (Figure 3). First, for a better fit with the S1’ pocket, we optimized the P1’ ligand while keeping the hydrogen bond with Thr26. As a result, compound **2**, having 6-chloro-2-methyl-2*H*-indazole as a P1’ ligand, displayed a 90-fold improvement in enzymatic inhibitory activity while maintaining the favorable DMPK profile. Next, the P1 methyl-amide moiety was replaced with a range of heterocyclic compounds, thus yielding compound **3**, which eventually became the clinical candidate, S-217622. S-217622 showed a biochemical activity of IC_50_ = 0.013 μM, an antiviral activity of EC_50_ = 0.37 μM, and preferable DMPK profiles for oral dosing, such as high metabolic stability (96% and 88% in human and rat liver microsomes, respectively), high oral absorption (97%) and low clearance (1.70 mL/min/mg) in rats (Figure 3, Tables S1–S3). Furthermore, S-217622 showed even better DMPK profiles in monkeys and dogs than in rats, with low clearance, long half-lives (t_1/2_) of approximately 10 and 30 hours in monkeys and dogs, respectively, and high oral bioavailability for all animals tested, suggesting its potential use for once-daily treatment of COVID-19 without requiring a PK booster such as ritonavir.

Figure 5 shows the X-ray co-crystal structure of 3CL^pro^ complexed with S-217622. In the S1 site, the 1-methyl-1*H*-1,2,4-triazole unit fit to the S1 pocket, forming a hydrogen bond with the side-chain NH of His163. The distinctive His41 flip observed in **1** was maintained in the S-217622 complex, and the 2,4,5-trifluorobenzylic moiety occupied the hydrophobic S2 pocket and stacked with the side chain of His41. The P1’ ligand, 6-chloro-2-methyl-2*H*-indazole moiety held hydrogen bonding with the Thr26 main-chain NH and hydrophobic contact with Met49 as seen in the co-crystal structure of **1**.

**Figure 5.**
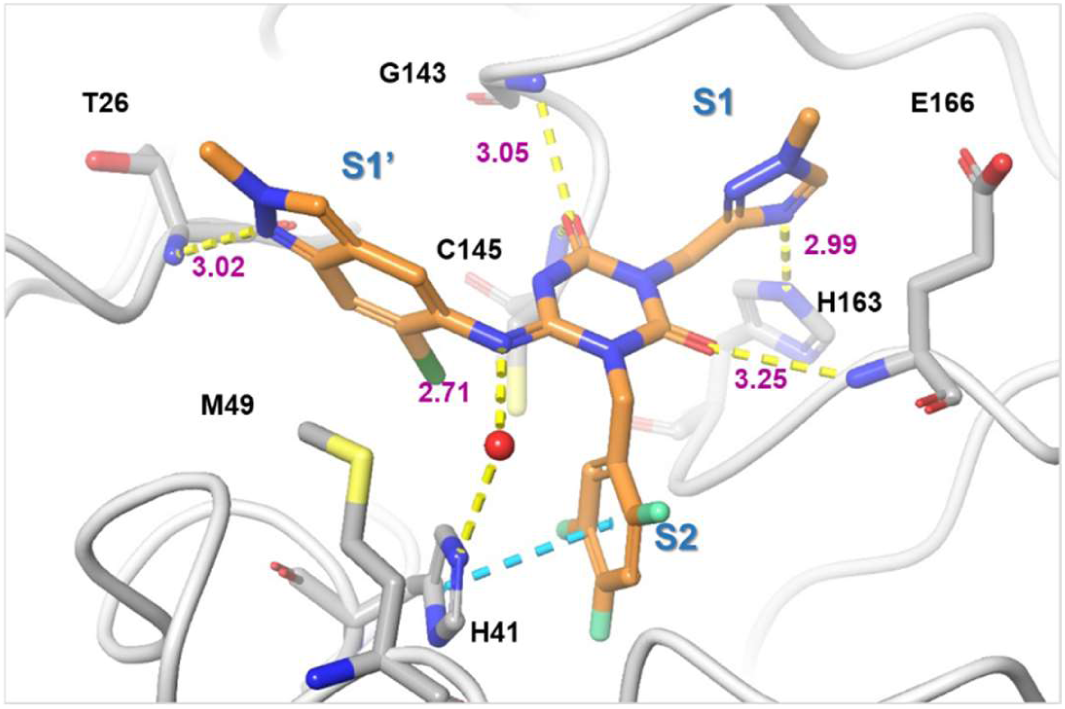
X-ray co-structure of S-217622 (**3**) and 3CL^pro^. **3** is colored in orange and the protein is colored in gray. Water molecules are shown as red spheres. Hydrogen bonds are indicated as yellow dashed lines; π-π stacking is indicated as a cyan dashed line.

Figure 6 summarizes the *in vitro* antiviral activities of S-217622 against several clinically isolated SARS-CoV-2 variants and family of coronaviruses. The antiviral activities were evaluated as per their inhibitory ability of the cytopathic effects elicited in SARS-CoV-2-infected VeroE6/TMPRSS2 cells. S-217622 exhibited similar antiviral activities against all tested SARS-CoV-2 variants, including the omicron strain, which has caused the current wave of the pandemic, indicating its potential broad usability as a therapeutic agent for treating COVID-19 (half-maximal effective concentration [EC_50_]: 0.29–0.50 μM; Figure 6a, Tables S2, S3). Because no significant mutations have been reported near the catalytic center of 3CL^pro^ in these variants of concern, orthosteric 3CL^pro^ inhibitors should be effective against all strains known to date. Antiviral activity of S-217622 against SARS-CoV (EC_50_: 0.21 μM, Figure 6b) was also comparable to that against SARS-CoV-2, where the sequence homology of 3CL^pro^ between SARS-CoV-2 and SARS-CoV was well conserved. S-217622 (**3**) also exhibited potent antiviral activity against MERS-CoV (EC_50_: 1.4 μM, Figure. 6c), HCoV-OC43 (EC_90_: 0.074 μM, Figure 6e) and HCoV-229E (EC_50_: 5.5 μM, Figure 6d). As described above, S-217622 displayed broad antiviral activities against a range of coronaviruses, suggesting possible applications of this compound or its derivatives for the next pandemic caused by future emerging coronaviruses. S-217622 showed no inhibitory activity against host-cell proteases, such as caspase-2, chymotrypsin, cathepsin B/D/G/L, and thrombin at up to 100 μM, suggesting its high selectivity for coronavirus proteases (Table 1). S-217622 (**3**) exhibited no safety concerns *in vitro* in studies involving ether-a-go-go-related gene inhibition, mutagenicity/clastogenicity, and phototoxicity (Table S4).

**Figure 6.**
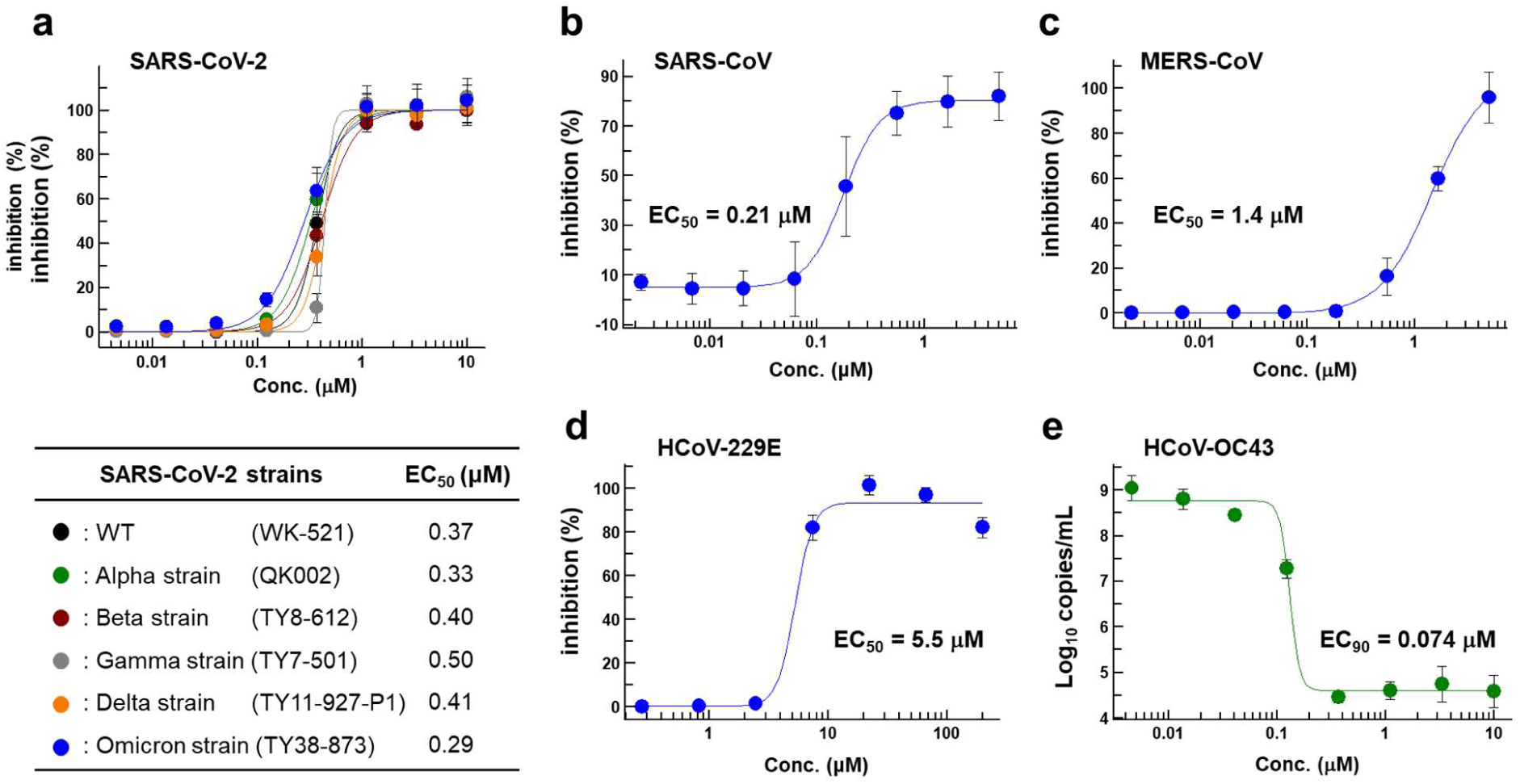
*In vitro* cellular activity of S-217622. Antiviral activity of S-217622 (fumaric acid co-crystal) against (a) various SARS-CoV-2 strains, (b) SARS-CoV, and (c) MERS-CoV in a cytopathic effect (CPE) inhibition assay using VeroE6/TMPRSS2 cells. Antiviral activity of S-217622 against (d) HCoV-229E (*Alphacoronavirus*) in a CPE inhibition assay with MRC-5 cells, and (e) HCoV-OC43 (*Betacoronavirus*) in a real-time quantitative reverse transcription polymerase chain reaction (RT-qPCR) assay with MRC-5 cells. Data are the means ± standard deviation; n = 3 biological replicates for SARS-CoV-2 strains, MERS-CoV, HCoV-229E, and HCoV-OC43 and n = 4 for SARS-CoV.

**Table 1.**
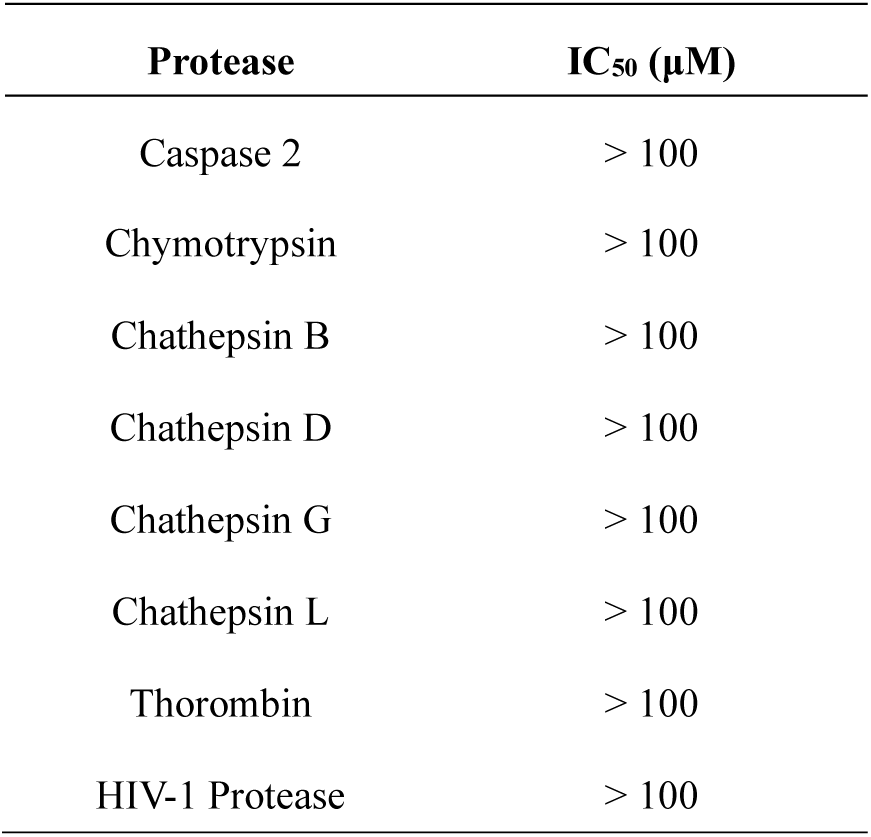
Enzymatic inhibitory activity of S-217622 against human host proteases and an HIV-1 protease. Inhibitory activities were < 50% at 100 μM.

We evaluated the antiviral efficacy of S-217622 *in vivo* in mice infected with SARS-CoV-2 Gamma strain (Figure 7). K417T, E484K, and N501Y mutations at the receptor-binding domain of the spike protein in SARS-CoV-2 Gamma strain promote interactions with mouse ACE2.^20^ Five-week-old BALB/c mice were intranasally inoculated with SARS-CoV-2 Gamma strain (hCoV-19/Japan/TY7-501/2021), and S-217622 (**3**) was administered orally as a 0.5% methylcellulose suspension immediately and twelve hours after infection (Figure 7a). Twenty-four hours after viral infection, the mice were euthanized, and the viral titers in their lung homogenates were measured. S-217622 (**3**) treatment reduced the intrapulmonary viral titers dose-dependently (Figure 7b). The mean viral titer was significantly lower in the S-217622 (**3**) treatment groups than in the vehicle treatment group (2 mg/kg vs vehicle, p = 0.0289; 8, 16 and 32 mg/kg vs vehicle, p < 0.0001). Viral titers reached near the lower limit of quantification (1.80-log_10_ 50% tissue culture infectious dose [TCID_50_]/mL) at 16 and 32 mg/kg in the S-217622 (**3**) treatment group. The plasma concentration increased dose-dependently between 2 and 32 mg/kg in the infected mice (Figure. 7c), and at doses of ≥16 mg/kg, the plasma concentration was estimated to be above the protein-adjusted-EC_50_ (PA-EC_50_) value (extrapolated to 100% mouse serum, 3.93 µmol/L = 2,090 ng/mL) over time, indicating the importance of the free plasma concentration for *in vivo* efficacy (Figure 7d). Although we applied twice-daily treatment in this mouse model, a once-daily treatment model could be applicable in clinical treatment because S-217622 showed a much lower clearance and longer half-lives in non-rodents than in rodents (Figure 3).

**Figure 7.**
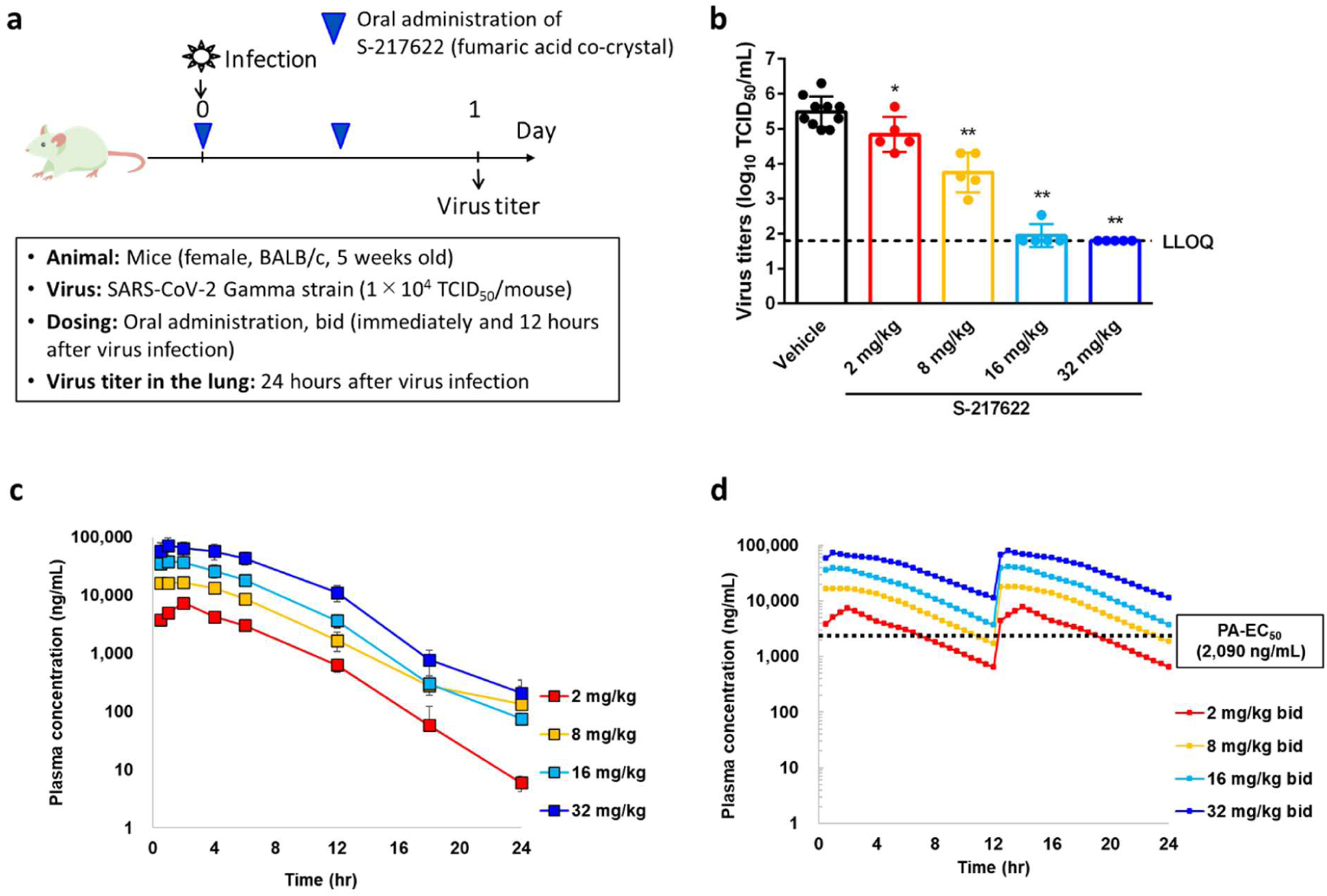
Dose-dependent *in vivo* antiviral efficacy of S-217622 in mice infected with SARS-CoV-2. (a) Protocol for the *in vivo* study. bid = twice a day; (b) Effect of S-217622 (fumaric acid co-crystal) treatment on lung viral titers in SARS-CoV-2 Gamma strain (hCoV-19/Japan/TY7-501/2021)-infected mice. TCID_50_ = 50% tissue culture infectious dose; each point represents an individual viral titer (n = 5–10). The broken line represents the lower limit of quantification (1.80 log10 TCID_50_/mL). The following *p*-values were calculated using Dunnett’s test: * *p* < 0.05 and ** *p* < 0.0001 vs vehicle. (c) S-217622 plasma concentration in the infected mice (n = 4). (d) Simulated S-217622 plasma concentrations after repeated oral administration of S-217622 (fumaric acid co-crystal) twice daily in infected mice as per non-parametric superposition. PA-EC_50_ = protein-adjusted EC_50_ extrapolated to 100% mouse serum.

## Chemistry

The synthetic scheme for compound **1** is described in Scheme 1. Starting from the pyrazole derivative **4**, cyclization with Ethyl isocyanatoacetate and CDI was conducted, giving **5** in 90% yield. Then, an alkylation with 5-bromomethyl-1,2,3-trifluorobenzene followed by introduction of a 4-difluoromethoxy-2-methylaniline unit, to give **7** (40% in 2 steps). The ester group in **7** was hydrolyzed and then amidated with methylamine, yielding **1** (58% in 2 steps). Compound **2** was synthesized similarly as shown in Scheme 2.

S-217622 (**3**) was synthesized as described in Scheme 3. Starting from known compound **9**,^21^ an alkylation with 1-(bromomethyl)-2,4,5-trifluorobenzene gave **10** in 93% yield. Then, the 3-*tert*-Bu group was removed and the triazole unit was introduced, and the substitution of the SEt moiety with the indazole unit finally gave S-217622 (**3**).

**Scheme 1.**
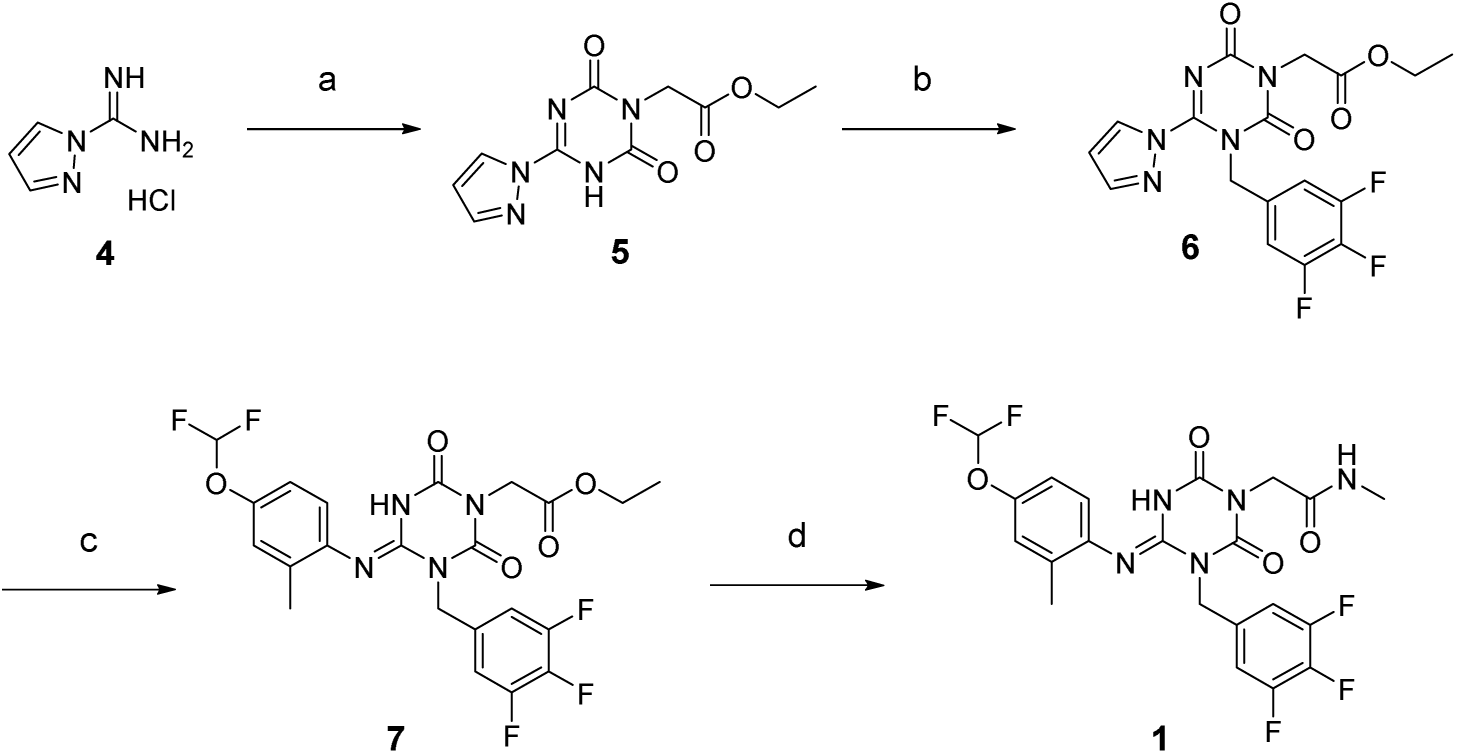
Reagents and Conditions: (a) ethyl isocyanato-acetate, DBU, CDI, DMA, –10 °C to rt, 90%; (b) 5-bromomethyl-1,2,3-trifluorobenzene, *N*,*N*-diisopropylethylamine, DMA, 60 °C; (c) 4-difluoromethoxy-2-methylaniline, *tert*-butanol, 100 °C, 40% in 2 steps; (d) (i) NaOH aq., THF/MeOH, rt; (ii) methylamine, HATU, *N*,*N*-diisopropylethylamine, THF, rt., 58% in 2 steps.

**Scheme 2.**
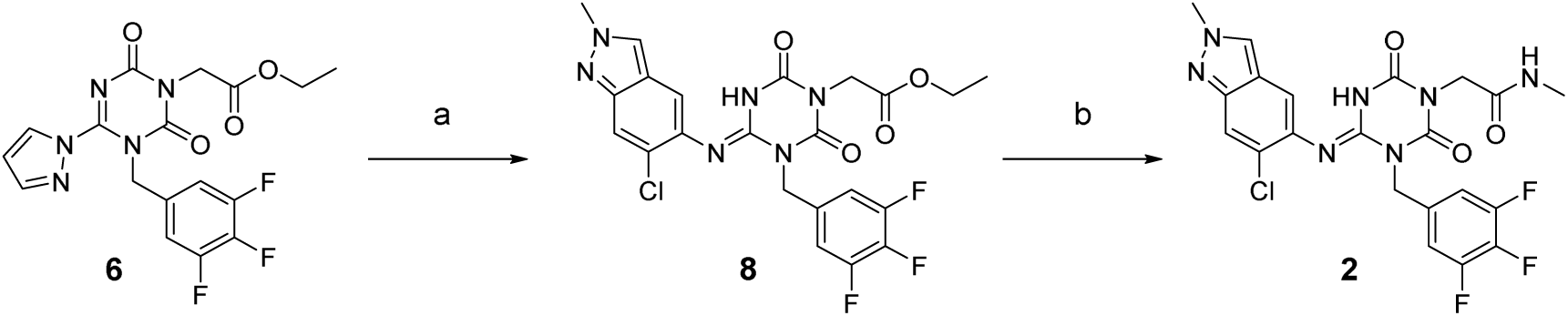
Reagents and Conditions: (a) 6-chloro-2-methyl-2*H*-indazol-5-amine, *tert*-amyl alcohol, 100 °C, 44% in 2 steps from **5**; (b) (i) NaOH aq., THF/MeOH, rt; (ii) methylamine, HATU, *N*,*N*-diisopropylethylamine, THF, rt., 29% in 2 steps.

**Scheme 3.**
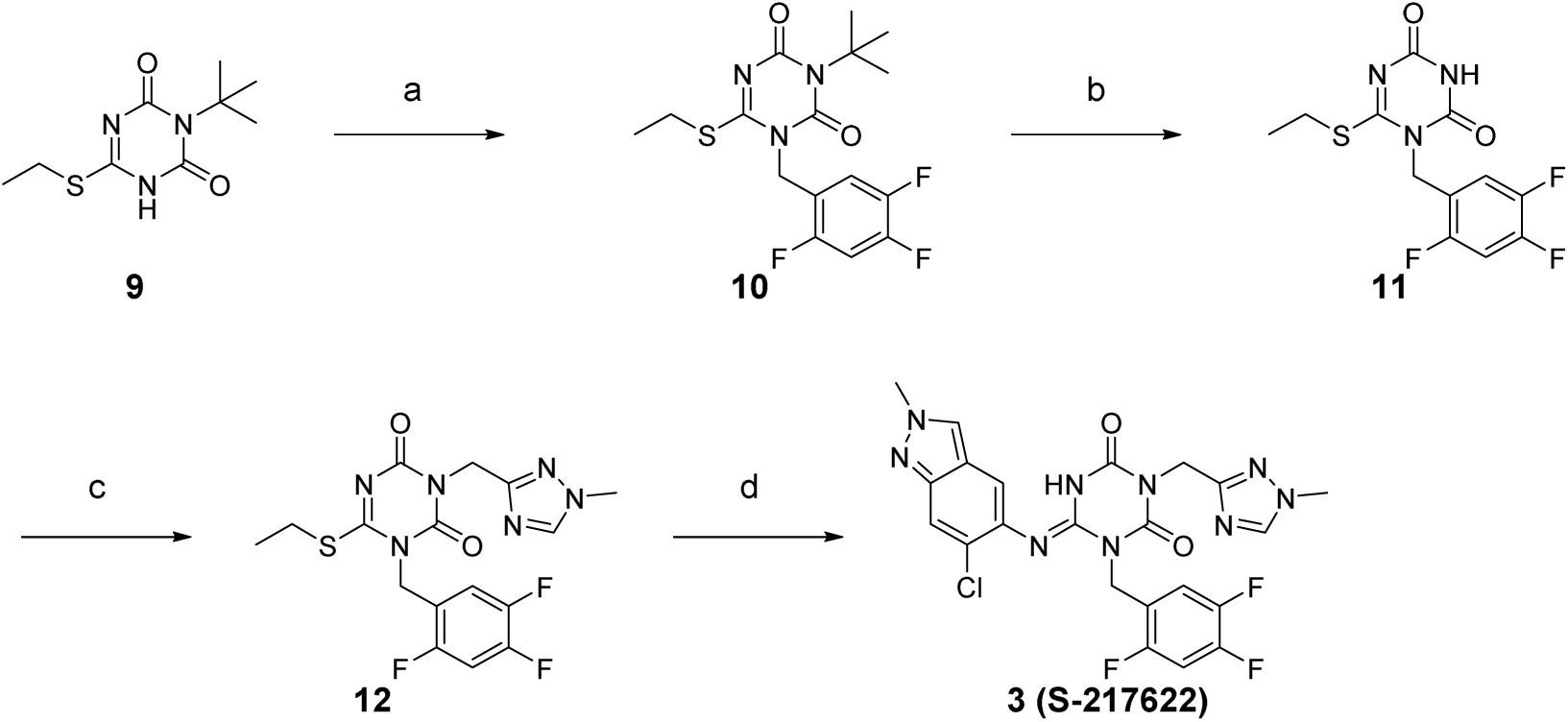
Reagents and Conditions: (a) 1-(bromomethyl)-2,4,5-trifluorobenzene, K_2_CO_3_, MeCN, 80 °C, 93%; (b) TFA, rt, 97%; (c) 3-(chloromethyl)-1-methyl-1*H*-1,2,4-triazole hydrochloride, K_2_CO_3_, DMF, 60 °C, 45%; (d) 6-chloro-2-methyl-2*H*-indazol-5-amine, LHMDS, THF, 0 °C to rt., 25%.

## Conclusions

Here, we described the discovery of S-217622, the first non-peptidic, non-covalent, oral 3CL^pro^ inhibitor clinical candidate for treating COVID-19. When we started this discovery program, most of the known inhibitors were peptide substrate mimetics with covalent warheads that bound covalently to Cys145 in the active site of 3CL^pro^. We assumed that these peptidic and reactive structural features would cause problems in the DMPK profile, such as low oral bioavailability due to low cell permeability, low metabolic stability, and low stability in the blood serum. Thus, we began the *de novo* search for non-peptidic 3CL^pro^ inhibitors using the SBDD strategy to combat the current SARS-CoV-2 pandemic. Virtual screening followed by biological screening yielded several hit compounds with IC_50_ values <10 µM, and one of these hit compounds, compound **1**, showed a favorable DMPK profile for an oral agent. Using the X-ray co-structure, SBDD-based structural optimization enabled >600-fold activity improvement while maintaining a good DMPK profile; this ultimately yielded the drug candidate, S-217622 (**3**). S-217622 (**3**) exhibited a favorable preclinical profile as a once-daily oral therapeutic agent for COVID-19 with promising antiviral activities to known variants of concern, a long half-life *in vivo*, especially in monkeys and dogs, excellent oral bioavailability, and steep efficacy in an *in vivo* mouse model infected with SARS-CoV-2. These favorable profiles prompted us to progress S-217622 to clinical trials, and studies are ongoing.

## Experimental section

### General Chemistry

All commercial reagents and solvents were used as received without further purification. Reactions were monitored *via* thin-layer chromatography performed on Merck silica gel plates (60 F254) or analytical liquid chromatography/mass spectroscopy (LC/MS) performed on a Shimadzu Shim-pack XR-ODS (C18, 2.2 µm, 3.0 × 50 mm, linear gradient from 10% to 100% B over 3 min, then 100% B for 1 min [A = water + 0.1% formic acid, B = MeCN + 0.1% formic acid], flow rate: 1.6 mL/min) using a Shimadzu UFLC system equipped with a LCMS-2020 mass spectrometer, LC-20AD binary gradient module, SPD-M20A photodiode array detector (detection at 254 nm), and SIL-20AC sample manager. All compounds used in the bioassay are >95% pure by HPLC analysis. Flash column chromatography was performed on an automated purification system using Fuji Silysia prepacked silica gel columns. ^1^H and ^13^C NMR spectra were recorded on a Bruker Advance at 400 and 100 MHz, respectively. Spectral data are reported as follows: chemical shift (as ppm referenced to tetramethylsilane), integration value, multiplicity (s = singlet, d = doublet, t = triplet, q = quartet, m = multiplet, br = broad), and coupling constant. High-resolution mass spectra were recorded on a Thermo Fisher Scientific LTQ Orbitrap using electrospray positive ionization.

#### Ethyl 2-[2,6-dioxo-4-(1H-pyrazol-1-yl)-3,6-dihydro-1,3,5-triazin-1(2H)-yl]acetate (5)

To a stirred solution of 1*H*-pyrazole-1-carboximidamide hydrochloride **4** (53.5 g, 365 mmol) in DMA (214 mL) were added ethyl isocyanatoacetate (49.5 g, 383 mmol) and DBU (57.8 mL, 383 mmol) below –10 °C. The reaction mixture was allowed to warm to 0 °C and stirred for 30 min at the same temperature. CDI (89.0 g, 548 mmol) and DBU (85.0 mL, 566 mmol) were added to the mixture below 10 °C. After stirring at room temperature overnight, the reaction mixture was quenched with aqueous 2M HCl (1000 mL). The solid was filtered, washed with H_2_O to afford **5** (86.8 g, 90%) as a white solid. ^1^H NMR (400 MHz, DMSO-*d*_6_) δ 1.21 (3H, t, *J* = 7.1 Hz), 4.15 (2H, q, *J* = 7.1 Hz), 4.52 (2H, s), 6.74 (1H, dd, *J* = 2.9, 1.5 Hz), 8.08 (1H, d, *J* = 1.0 Hz), 8.59 (1H, dd, *J* = 2.9 Hz). ^13^C NMR (100 MHz, DMSO-*d*_6_) δ 14.03, 42.19, 61.24, 111.38, 130.47, 145.82, 151.32, 152.12, 167.64. HRMS-ESI (m/z): [M + H]^+^ calcd for [C_10_H_12_N_5_O_4_]^+^ 266.0877; found 266.0884.

#### Ethyl (4E)-2-(4-{[4-(difluoromethoxy)-2-methylphenyl]imino}-2,6-dioxo-3-(3,4,5-trifluorobenzyl)-1,3,5-triazinan-1-yl)acetate (7)

A mixture of **5** (1.06 g, 4.00 mmol), *N*,*N*-diisopropylethylamine (0.907 mL, 5.20 mmol), and 5-bromoethyl-1,2,3-trifluorobenzene (0.631 mL, 4.80 mmol) in DMA (10 mL) was stirred at 60 °C for 5 h. Then, the reaction mixture was cooled to room temperature, and diluted with H_2_O and EtOAc. The aqueous layer was extracted with EtOAc. The organic layer was washed with H_2_O and brine, dried over MgSO_4_, and concentrated under reduced pressure to afford a crude product of **6** (1.65 g). After a mixture of **6** (268 mg, ≤0.665 mmol) and 4-(difluoromethoxy)-2-methylaniline (0.094 mL, 0.665 mmol) in *tert*-butanol (2.7 mL) was stirred at 100 °C for 30 min, the reaction mixture was allowed to cool to room temperature. The mixture was concentrated under reduced pressure. The residue was purified by silica gel column chromatography (*n*-hexane/EtOAc, gradient, 15-33% EtOAc) to afford **7** (134 mg, 40% over 2 steps) as a white solid. ^1^H NMR (400 MHz, CDCl_3_) δ 1.31 (3H, t, *J* = 7.2 Hz), 2.01 (3H, s), 4.26 (2H, q, *J* = 7.2 Hz), 4.57 (2H, s), 5.17 (2H, s), 6.48 (1H, t, *J* = 74.0 Hz), 6.72 (1H, d, *J* = 8.5 Hz), 6.98 (1H, dd, *J* = 8.5, 2.4 Hz), 7.03 (1H, d, *J* = 2.4 Hz), 7.19 (2H, t, *J* = 7.3 Hz), 7.40 (1H, br s). ^13^C NMR (100 MHz, CDCl_3_) δ 14.11, 17.92, 42.71, 44.95, 62.16, 113.36 (dd, *J* = 16.1, 5.9 Hz), 115.95 (t, *J* = 260 Hz), 118.88, 121.78, 122.97, 132.09, 132.03-132.23 (m), 136.80, 139.52 (dt, *J* = 251.9, 15.2 Hz), 140.50, 147.13, 147.80 (t, *J* = 2.9 Hz), 149.46, 151.07 (ddd, *J* = 250.2, 9.5, 3.7 Hz), 167.35. HRMS-ESI (m/z): [M + H]^+^ calcd for [C_22_H_20_F_5_N_4_O_5_]^+^ 515.1359; found 515.1348.

#### (4E)-2-(4-{[4-(difluoromethoxy)-2-methylphenyl]imino}-2,6-dioxo-3-(3,4,5-trifluorobenzyl)-1,3,5-triazinan-1-yl)-N-methylacetamide (1)

A stirred solution of **7** (100 mg, 0.194 mmol) in THF/MeOH (2 mL, v/v = 1/1) was added NaOH (1M aqueous solution, 1.37 mL, 1.37 mmol) at room temperature. After stirring at room temperature for 2 h, the reaction mixture was quenched with 1 M aqueous HCl solution. The aqueous layer was extracted with EtOAc. The organic layer was washed with H_2_O and brine, dried over MgSO_4_, and concentrated under reduced pressure to afford a crude residue (71.1 mg). The residue (71.1 mg, ≤0.146 mmol) in THF (0.6 mL) were added methylamine (2M in THF, 0.110 mL, 0.219 mmol), *N*,*N*-diisopropylethylamine (0.077 mL, 0.439 mmol), and HATU (83.0 mg, 0.219 mmol). After stirring at room temperature overnight, the reaction mixture was diluted with H_2_O and EtOAc. The aqueous layer was extracted with EtOAc. The organic layer was washed with H_2_O and brine, dried over MgSO_4_, and concentrated under reduced pressure. The residue was purified by silica gel column chromatography (CHCl_3_/MeOH, gradient, 0-10% MeOH). The collected fraction was recrystallized from *n*-hexane/EtOAc to afford **1** (60.7 mg, 58% over 2 steps) as a white solid. ^1^H NMR (400 MHz, CDCl_3_) δ 1.97 (3H, s), 2.86 (3H, d, *J* = 4.8 Hz), 4.33 (2H, s), 5.13 (2H, s), 5.76 (1H, d, *J* = 4.8 Hz), 6.47 (1H, t, *J* = 74.2 Hz), 6.68 (1H, d, *J* = 8.3 Hz), 6.92 (1H, dd, *J* = 8.3, 2.4 Hz), 6.97 (1H, d, *J* = 2.4 Hz), 7.16 (2H, t, *J* = 7.4 Hz), 8.15 (1H, br s). ^13^C NMR (100 MHz, CDCl_3_) δ 17.93, 26.49, 43.67, 44.96, 113.27 (dd, *J* = 16.1, 5.9 Hz), 116.18 (t, *J* = 259.7 Hz), 118.59, 122.03, 122.56, 122.9, 132.21, 132.21-132.39 (m), 136.83, 139.44 (dt, *J* = 252.4, 15.0 Hz), 140.87, 147.56 (t, *J* = 2.9 Hz), 147.84, 149.68, 151.02 (ddd, *J* = 250.2, 10.3, 3.7 Hz), 166.40. HRMS-ESI (m/z): [M + H]^+^ calcd for [C_21_H_19_F_5_N_5_O_4_]^+^ 500.1352; found 500.1348.

#### Ethyl (4E)-2-{4-[(6-chloro-2-methyl-2H-indazol-5-yl)imino]-2,6-dioxo-3-(3,4,5-trifluorobenzyl)-1,3,5-triazinan-1-yl}acetate (8)

A mixture of **6** (291 mg, ≤ 0.711 mmol) and 6-chloro-2-methyl-2*H*-indazol-5-amine^22^ (129 mg, 0.711 mmol) in *tert*-amyl alcohol (3 mL) was stirred at 100 °C for 2 h. The reaction mixture was cooled to room temperature, and then concentrated under reduced pressure. The residue was triturated with EtOAc and the solid was filtered, washed with EtOAc to afford **8** (160 mg, 44% over 2 steps) as a white solid. ^1^H NMR (400 MHz, DMSO-*d*_6_, DCl in D_2_O) δ 1.18 (3H, t, *J* = 7.2 Hz), 4.12 (2H, q, *J* = 7.2 Hz), 4.15 (3H, s), 4.47 (2H, s), 5.28 (2H, s), 7.37 (2H, dd, *J* = 8.8, 7.0 Hz), 7.47 (1H, s), 7.74 (1H, s), 8.37 (1H, s). ^13^C NMR (100 MHz, DMSO-*d_6_*, DCl in D_2_O) δ 14.04, 40.20, 42.78, 44.44, 61.25, 79.33, 111.93 (dd, *J* = 16.1, 5.1 Hz), 116.84, 120.66, 125.43, 127.92, 128.65, 131.63, 133.46-133.65 (m), 137.98 (dt, *J* = 248.2, 15.4 Hz), 146.39, 150.22 (ddd, *J* = 247.0, 10.1, 3.9 Hz), 150.39, 150.52, 168.00. HRMS-ESI (m/z): [M + H]^+^ calcd for [C_22_H_19_ClF_3_N_6_O_4_]^+^ 523.1103; found 523.1104.

#### (4E)-2-{4-[(6-Chloro-2-methyl-2H-indazol-5-yl)imino]-2,6-dioxo-3-(3,4,5-trifluorobenzyl)-1,3,5-triazinan-1-yl)-N-methylacetamide (2)

To a stirred solution of **8** (445 mg, 0.851 mmol) in THF/MeOH (4.5 mL, v/v = 1/1) was added aqueous NaOH (1M solution, 2.55 mL, 2.55 mmol) at room temperature. After stirring at room temperature for 70 min, the reaction mixture was diluted with EtOAc and then quenched with aqueous 2 M HCl solution. The aqueous layer was extracted with EtOAc. The organic layer was washed with H_2_O and brine, dried over Na_2_SO_4_, and concentrated under reduced pressure. The residue was triturated with diisopropylether and the solid was filtered, washed with diisopropylether to afford a crude residue (467 mg). The crude residue (100 mg, ≤0.202 mmol) in THF (2 mL) were added methylamine (2M in THF, 0.152 mL, 0.303 mmol), *N*,*N*-diisopropylethylamine (0.106 mL, 0.606 mmol), and HATU (115 mg, 0.303 mmol). After stirring at room temperature overnight, the reaction mixture was diluted with H_2_O and EtOAc. The aqueous layer was extracted with EtOAc. The organic layer was washed with H_2_O and brine, dried over MgSO_4_, and concentrated under reduced pressure. The residue was purified by silica gel column chromatography (CHCl_3_/MeOH, gradient, 0-2% MeOH). The collected fraction was recrystallized from *n*-hexane/EtOAc to afford **2** (26.6 mg, 29% over 2 steps) as a white solid. ^1^H NMR (400 MHz, DMSO-*d*_6_, DCl in D_2_O) δ 2.57 (3H, s), 4.15 (3H, s), 4.27 (2H, s), 5.24 (2H, s), 7.38 (1H, s), 7.41 (2H, dd, *J* = 8.7, 7.2 Hz), 7.73 (1H, s), 8.36 (1H, s). ^13^C NMR (100 MHz, DMSO-*d*_6_, DCl in D_2_O) δ 25.47, 40.19, 43.99, 44.47, 79.37, 111.98 (dd, *J* = 16.9, 5.1 Hz), 115.78, 116.77, 120.81, 125.39, 128.64, 132.87, 133.78-133.97 (m), 137.91 (dt, *J* = 248.0, 15.4 Hz), 145.57, 146.14, 150.22 (ddd, *J* = 246.7, 9.9, 4.0 Hz), 150.30, 150.51, 166.97. HRMS-ESI (m/z): [M + H]^+^ calcd for [C_21_H_18_ClF_3_N_7_O_3_]^+^ 508.1106; found 508.1106.

#### 3-(tert-Butyl)-6-(ethylthio)-1,3,5-triazine-2,4(1H,3H)-dione (9)

Compound **9** was prepared according to the reported procedure.^21^ ^1^H NMR (400 MHz, CDCl_3_) δ 1.36 (3H, t, *J* = 7.4 Hz), 1.66 (9H, s), 3.14 (2H, q, *J* = 7.4 Hz). ^13^C NMR (100 MHz, CDCl_3_) δ 14.35, 25.22, 29.20, 61.55, 152.26, 154.19, 165.96. HRMS-ESI (m/z): [M + H]^+^ calcd for [C_9_H_16_N_3_O_2_S]^+^ 230.0958; found 230.0952.

#### 3-(tert-Butyl)-6-(ethylthio)-1-(2,4,5-trifluorobenzyl)-1,3,5-triazine-2,4(1H,3H)-dione (10)

A mixture of **9** (100 mg, 0.436 mmol), potassium carbonate (78.0 mg, 0.567 mmol), and 1-(bromomethyl)-2,4,5-trifluorobenzene (0.063 mL, 0.480 mmol) in MeCN (0.8 mL) was stirred at 80 °C for 2 h. The reaction mixture was cooled to room temperature, and then the mixture was diluted with EtOAc. The precipitate was filtered, and the filtrate was concentrated under reduced pressure. The residue was purified by silica gel column chromatography (*n*-hexane/EtOAc gradient, 0-30% EtOAc) to afford **10** (151 mg, 93%) as a colorless oil. ^1^H NMR (400 MHz, CDCl_3_) δ 1.33 (3H, t, *J* = 7.4 Hz), 1.65 (9H, s), 3.15 (2H, q, *J* = 7.4 Hz), 5.03 (2H, s), 6.91-7.01 (2H, m). ^13^C NMR (100 MHz, CDCl_3_) δ 13.73, 26.88, 28.88, 41.05 (d, *J* = 4.4 Hz), 61.59, 105.92 (dd, *J* = 27.5, 20.9 Hz), 116.24 (ddd, *J* = 20.5, 5.1, 1.5 Hz), 118.65 (td, *J* = 10.5, 5.4 Hz), 147.03 (ddd, *J* = 246.1, 12.8, 3.7 Hz), 149.72 (ddd, *J* = 252.4, 13.9, 12.5 Hz), 150.36, 152.98, 155.24 (ddd, *J* = 246.5, 9.5, 2.9 Hz), 166.69. HRMS-ESI (m/z): [M + H]^+^ calcd for [C_16_H_19_F_3_N_3_O_2_S]^+^ 374.1145; found 374.1142.

#### 6-(Ethylthio)-1-(2,4,5-trifluorobenzyl)-1,3,5-triazine-2,4(1H,3H)-dione (11)

A mixture of **10** (4.88 g, 13.08 mmol) in TFA (9.8 mL) was stirred at room temperature for 4 h then stood at the same temperature overnight. After concentration under reduced pressure, the residue was azeotroped with toluene and triturated with diisopropylether to afford **11** (4.01 g, 97%) as a white solid. ^1^H NMR (400 MHz, CDCl_3_) δ 1.37 (3H, t, *J* = 7.4 Hz), 3.23 (2H, q, *J* = 7.4 Hz), 5.15 (2H, s), 6.95-7.09 (2H, m), 8.23 (1H, br s). ^13^C NMR (100 MHz, DMSO-*d*_6_) δ 13.73, 26.40, 40.66 (d, *J* = 3.7 Hz), 106.13 (dd, *J* = 28.2, 21.6 Hz), 116.51 (dd, *J* = 20.9, 4.8 Hz), 119.52 (dq, *J* = 16.1, 3.2 Hz), 145.05-149.96, 150.08, 152.42, 154.70 (ddd, *J* = 244.5, 10.1, 2.4 Hz), 169.98. HRMS-ESI (m/z): [M + H]^+^ calcd for [C_12_H_11_FN_3_O_2_S]^+^ 318.0519; found 318.0516.

#### 6-(Ethylthio)-3-[(1-methyl-1H-1,2,4-triazol-3-yl)methyl]-1-(2,4,5-trifluorobenzyl)-1,3,5-triazine-2,4(1H,3H)-dione (12)

A mixture of **11** (2.50 g, 7.88 mmol), 3-(chloromethyl)-1-methyl-1*H*-1,2,4-triazole hydrochloride (1.99 g, 11.8 mmol), and potassium carbonate (3.27 g, 23.6 mmol) in DMF (23 mL) was stirred at 60 °C for 3.5 h. The reaction mixture was allowed to cool to room temperature and diluted with aqueous NH_4_Cl solution. The precipitate was filtered and washed with H_2_O. The solid was purified by silica gel column chromatography (*n*-hexane/EtOAc gradient, 30-60% EtOAc) to afford **12** (1.47 g, 45%) as a white solid. ^1^H NMR (400 MHz, CDCl_3_) δ 1.34 (3H, t, *J* = 7.4 Hz), 3.20 (2H, q, *J* = 7.4 Hz), 3.84 (3H, s), 5.16 (2H, s), 5.23 (2H, s), 6.92-6.98 (1H, m), 7.10-7.17 (1H, m), 7.93 (1H, s). ^13^C NMR (100 MHz, CDCl_3_) δ 13.55, 27.31, 36.13, 39.92, 41.11 (d, *J* = 3.7 Hz), 105.80 (dd, *J* = 27.5, 20.9 Hz), 116.20 (dd, *J* = 20.5, 3.7 Hz), 118.11 (td, *J* = 11.0, 4.9 Hz), 144.33, 145.87-151.14, 150.57, 151.68, 155.18 (ddd, *J* = 246.5, 9.5, 2.9 Hz), 159.29, 169.56. HRMS-ESI (m/z): [M + H]^+^ calcd for [C_16_H_16_F_3_N_6_O_2_S]^+^ 413.1002; found 413.0998.

#### (6E)-6-[(6-Chloro-2-methyl-2H-indazol-5-yl)imino]-3-[(1-methyl-1H-1,2,4-triazol-3-yl)methyl]-1-(2,4,5-trifluorobenzyl)-1,3,5-triazinane-2,4-dione (3, S-217622)

To a solution of **12** (300 mg, 0.727 mmol) and 6-chloro-2-methyl-2*H*-indazol-5-amine (172 mg, 0.946 mmol) in THF (6 mL) was added LHMDS (1M in THF; 1.46 mL, 1.46 mmol) dropwisely at 0 °C. The reaction mixture was stirred at 0 °C for 2.5 h and then at rt for 40 min. The reaction was quenched with aqueous NH_4_Cl solution, and the aqueous layer was extracted with EtOAc. The organic layer was washed with brine, dried over MgSO_4_, and concentrated under reduced pressure. The residue was purified by silica gel column chromatography (CHCl_3_/MeOH gradient, 0-20% MeOH). The solid was recrystallized from acetone/H_2_O to afford **3 (S-217622)** (95.3 mg, 25%) as a pale brown solid. ^1^H NMR (400 MHz, DMSO-*d*_6_, DCl in D_2_O) δ 3.90 (3H, s), 4.15 (3H, s), 5.04 (2H, s), 5.26 (2H, s), 7.44 (1H, m), 7.52-7.65 (2H, m), 7.73 (1H, s), 8.40 (1H, s), 9.31 (1H, s). ^13^C NMR (100 MHz, DMSO-*d*_6_, DCl in D_2_O) δ 37.34, 38.04, 40.06, 40.29, 106.16 (dd, *J* = 28.2, 21.6 Hz), 116.46-116.70, 116.70, 120.54-120.76, 120.76, 125.93, 129.10, 132.35, 143.84, 145.98, 146.38 (ddd, *J* = 241.4, 12.5, 3.7 Hz), 146.60, 148.52 (td, *J* = 247.7, 13.6 Hz), 150.43, 150.50, 155.22 (ddd, *J* = 244.3, 10.3, 2.2 Hz), 155.58. HRMS-ESI (m/z): [M + H]^+^ calcd for [C_22_H_18_ F_3_ClN_9_O_2_]^+^ 532.1219; found 532.1221.

### Preparation of Compound 3 (S-217622) fumaric acid co-crystal

A mixture of **3 (S-217622)** (1.17 g, 2.2 mmol) and fumaric acid (278 mg, 2.4 mmol) in EtOAc (5.9 mL) was stirred at room temperature for 45 min. The suspension was filtrated to afford **3** (S-217622) **fumaric acid co-crystal** (1.37 g, 95 %) as a white solid. ^1^H NMR (400 MHz, pyridine-*d_5_*) δ 3.64 (s, 3H), 3.99 (s, 3H), 5.56 (s, 2H), 5.61 (s, 2H), 7.16-7.25 (m, 2H), 7.44 (s, 2H), 7.81 (s, 1H), 7.89 (s, 1H), 7.89-7.97 (m, 1H), 8.32 (s, 1H).

#### Cells and viruses

Remdesivir was purchased from MedChemExpress. VeroE6/TMPRSS2 cells from the National Institutes of Biomedical Innovation (Tokyo, Japan) were used to evaluate the antiviral activity against SARS-CoV-2. Those prepared by Hokkaido University as previously reported^23^ were used to evaluate the antiviral activities against SARS-CoV and MERS-CoV. MRC-5 cells (CCL-171) were purchased from American Type Culture Collection (ATCC; Manassas, VA, USA). Cells were maintained in Dulbecco’s modified Eagle’s medium (Thermo Fisher Scientific) supplemented with 10% heat-inactivated fetal bovine serum (FBS) (Sigma-Aldrich Co., Ltd.) at 37°C with 5% CO_2_.

SARS-CoV-2 clinical isolates were obtained from the National Institute of Infectious Diseases (NIID; Tokyo, Japan): hCoV-19/Japan/TY/WK-521/2020 (Pango Lineage: A), hCoV-19/Japan/QK002/2020 (Alpha, B.1.1.7), hCoV-19/Japan/QHN001/2020 (Alpha, B.1.1.7), hCoV-19/Japan/QHN002/2020 (Alpha, B.1.1.7), hCoV-19/Japan/TY7-501/2021 (Gamma, P.1), hCoV-19/Japan/TY7-503/2021 (Gamma, P.1), hCoV-19/Japan/TY8-612/2021 (Beta, B.1.351), hCoV-19/Japan/TY11-927-P1/2021 (Delta, B.1.617.2), and hCoV-19/Japan/TY38-873/2021 (Omicron, B.1.1.529). All SARS-CoV-2 strains were propagated in VeroE6/TMPRSS2 cells, and infectious titers were determined by standard tissue culture infectious dose (TCID)_50_ in VeroE6/TMPRSS2 cells. SARS-CoV (Hanoi strain) was provided by Dr. Koichi Morita of Nagasaki University^24^. MERS-CoV (EMC/2012) was provided by Dr. Bart L Haagmans, Erasmus University Medical Center^25^. VeroE6 cells (ATCC) were used to propagate SARS-CoV; VeroE6/TMPRSS2 cells were used to propagate MERS-CoV. HCoV-OC43 and HCoV-229E were obtained from ATCC.

#### 3CL protease inhibition assay

The 3CL protease inhibition assay was conducted in 384-well plates (Corning 3702). The substance solution (10 mM dimethyl sulfoxide [DMSO] solution) was diluted to 250 µmol/L stepwise with a 3-fold dilution with DMSO. Finally, the solutions were mixed with 20 mmol/L Tris-HCl (pH 7.5) as a compound solution. Ten microliters of compound solution was added manually to each well, then 5 µL of 16 µM substrate in inhibition buffer (2 mM EDTA, 20 mM DTT, 0.02% BSA, and 20 mM Tris-HCl, pH 7.5) was added. The reaction was initiated by adding 5 µL of 12 nM 3CL protease in inhibition buffer and incubated at room temperature for 3 h. The following operations were the same as those described in the biological screening.

#### Biological Screening

The compound screening assay was performed in 384-well plates (Corning 3702 or Greiner 781280). One-hundred fifty nanoliters of testing compound at various concentrations was added to each well by an ECHO 555 dispenser (LABCYTE INC.). Next, 7.5 µL of 8 µM substrate (Dabcyl-KTSAVLQSGFRKME [Edans] -NH2, 3249-v, PEPTIDE INSTITUTE, Inc.) in assay buffer (100 mM NaCl, 1 mM ethylenediaminetetraacetic acid [EDTA], 10 mM DL-dithiothreitol (DTT), 0.01% bovine serum albumin [BSA], and 20 mM Tris-HCl, pH 7.5) was dispensed using Multidrop Combi (Thermo Scientific). The reaction was initiated by adding 7.5 µL of 6 or 0.6 nM 3CL protease in assay buffer and incubated at room temperature for 3 h. After incubation, the reaction was stopped by adding 45 µL of water solution containing 0.1% formic acid, 10% acetonitrile, and 0.05 µmol/L Internal Standard (IS) peptide (Dabcyl-KTSAVLeu [^13^C_6_,^15^N]-Q, custom synthesized by PEPTIDE INSTITUTE, Inc.). The reactions were analyzed with MS using a RapidFire 360 high-throughput sampling robot (Agilent Technologies) connected to an iFunnel Agilent 6550 accurate mass quadrupole time-of-flight mass spectrometer using electrospray. Peak areas were acquired and analyzed using a RapidFire Integrator (Agilent Technologies). Reaction product peak areas were acquired from m/z 499.27; IS peak areas were acquired from m/z 502.78. IC_50_ values were determined by plotting compound concentration vs inhibition and fitting data with a 4-parameter logistical fit (Model 205, XLfit).

#### Cellular Antiviral Activity

Antiviral activity against SARS-CoV-2, SARS-CoV, MERS-CoV and HCoV-229E was assessed by monitoring cell viability; that against HCoV-OC43 was assessed by monitoring viral RNA in a cell suspension. EC_50_ values were determined by plotting compound concentration vs inhibition and fitting data with a 4-parameter logistical fit (Model 205, XLfit). EC_90_ values against HCoV-OC43 were determined from the resulting dose-response curves and calculated with the two-point method.

Antiviral activities against SARS-CoV-2 were evaluated using VeroE6/TMPRSS2 cells. VeroE6/TMPRSS2 cells (1.5 × 10^4^/well) suspended in minimum essential medium (MEM) (Thermo Fisher Scientific) supplemented with heat-inactivated 2% FBS were seeded into 96-well plates with diluted compounds in each well. Cells were infected with each SARS-CoV-2 at 30–3000 TCID_50_/well and cultured at 37°C with 5% CO_2_ for 3 days or 4 days. Cell viability was assessed using a CellTiter-Glo® 2.0 assay (Promega). The CC_50_ was assessed in the absence of viruses after being cultured for 3 days.

Antiviral activities against SARS-CoV and MERS-CoV were evaluated at Hokkaido University using VeroE6/TMPRSS2 cells as previously reported^23^. VeroE6/TMPRSS2 cells (1.5 × 10^4^/well) suspended in 2% FBS-containing MEM were seeded into 96-well plates with diluted compounds in each well. Cells were infected with each SARS-CoV at 1000 TCID_50_/well or MERS-CoV 2500 TCID_50_/well and cultured at 37°C with 5% CO_2_ for 3 days. Cell viability was assessed via (3-[4,5-dimethyl-2-thiazolyl]-2,5-diphenyl-2*H*-tetrazolium bromide (MTT) assay (Nacalai Tesque) as previously described^26^.

Antiviral activity against HCoV-229E was evaluated using MRC-5 cells. MRC-5 cells (2.0 × 10^4^/well) suspended in 2% FBS-containing MEM were seeded into 96-well plates and incubated at 37 °C with 5% CO_2_ overnight. The next day, the cells were infected with HCoV-229E at 1000 TCID_50_/well and incubated at 37 °C with 5% CO_2_ for 1 h, followed by removal of the inoculum and added 2% FBS-containing MEM with the diluted compounds. Cells infected with HCoV-229E were incubated at 37 °C with 5% CO_2_ for 3 days. Cell viability was assessed using a CellTiter-Glo® 2.0 assay.

Antiviral activity against HCoV-OC43 was evaluated using MRC-5 cells. MRC-5 cells (2.0 × 10^4^/well) suspended in 2% FBS-containing MEM were seeded into 96-well plates and incubated at 37 °C with 5% CO_2_ overnight. The next day, the cells were infected with HCoV-OC43 at 100 TCID_50_/well and incubated at 37 °C with 5% CO_2_ for 1 h, followed by removal of the inoculum and added 2% FBS-containing MEM with the diluted compounds. Cells infected with HCoV-OC43 were incubated at 37 °C with 5% CO_2_ for 42 h, and viral RNA was extracted from the supernatants using a Quick-RNA Viral Kit (ZYMO RESEARCH, # R1041). Viral RNA was quantified via real-time PCR (Applied Biosystems, QuantStudio 3) with specific primers and probes for HCoV-OC43 detection^27^.

#### Cellular antiviral activity in the presence of mouse serum

Antiviral activity against SARS-CoV-2 in the presence of mouse serum was assessed by monitoring cell viability. S-217622 (fumaric acid co-crystal) was diluted with 3.125%, 6.25%, 12.5%, and 25% mouse serum in MEM supplemented with heat-inactivated 2% FBS. One hundred microliters of serially diluted compound solutions were added to a 96-well plate and incubated at room temperature for approximately 1 h. Each 50 µL/well of VeroE6/TMPRSS2 cells was adjusted to 3.0 × 10^5^ cells/mL with MEM supplemented with heat-inactivated 2% FBS and dispensed on the plate. Each 50 µL/well of SARS-CoV-2 was added at 10000 TCID_50_/well and cultured at 37 °C with 5% CO_2_ for 3 days. Cell viability was assessed using a CellTiter-Glo® 2.0 assay, followed by determination of the EC_50_ value from the cell viability. PA-EC_50_ extrapolated to 100% serum was calculated by linear regression using the EC_50_ value of each serum concentration. PS extrapolated to 100% serum was calculated by dividing the PA-EC_50_ (extrapolated value of 100% mouse serum) by EC_50_ (in the presence of mouse serum).

#### Human protease enzyme assay

Selectivity tests against varieties of host protease activity were conducted by Eurofins Panlabs Discovery Services Taiwan, Ltd. on behalf of SHIONOGI Co. & Ltd. as per established protocols. S-217622 (fumaric acid co-crystal) was tested on a set of seven proteases (caspase-2, chymotrypsin, cathepsin B/D/G/L and thrombin) at 100 µM.

#### *In vivo* SARS-CoV-2 infection and treatment studies

*In vivo* SARS-CoV-2 infection experiments were conducted in accordance with the guidelines of the Association for Assessment and Accreditation of Laboratory Animal Care (AAALAC). The animal study protocol was approved by the director of the institute based on the report of the Institutional Animal Care and Use Committee of Shionogi Research Laboratories.

Mouse i*n vivo* SARS-CoV-2 infection studies were done at Shionogi Pharmaceutical Research Center (Osaka, Japan). Five-week-old female BALB/cAJcl mice (CLEA Japan, Inc.; n=5 or 10 per group) were intranasally inoculated with SARS-CoV-2 Gamma strain (hCoV-19/Japan/TY7-501/2021) (10000 TCID_50_/mouse) under anesthesia. Immediately after infection, the mice were orally administered S-217622 (fumaric acid co-crystal) (2, 8, 16 or 32 mg/kg q12h; n=5 per group) or vehicle (0.5 w/v% methyl cellulose in aqueous solution q12h; n=10 per group), for 1 day. Twenty-four hours postinfection, the mice were euthanized via cervical dislocation under anesthesia; their lungs were removed and the viral titers in the lung homogenates were determined using VeroE6/TMPRSS2 cells. Viral titers are expressed as log_10_ TCID_50_/mL.

#### PK study in infected mice

PK experiments in infected mice were conducted in accordance with the guidelines provided by AAALAC and were approved by IACUC of Shionogi Research Laboratories.

Mouse PK studies were done at Shionogi Pharmaceutical Research Center (Osaka, Japan). BALB/cAJcl mice were intranasally inoculated with SARS-CoV-2 Gamma strain (hCoV-19/Japan/TY7-501/2021) (10000 TCID_50_/mouse) and orally administered with S-217622 (fumaric acid co-crystal) (2, 8, 16, or 32 mg/kg) immediately after infection. Blood was taken at 0.5, 1, 2, 4, 6, 12, 18 and 24 h after dosing (n = 4 per group per timepoint) and plasma concentrations of S-217622 were determined by LC/MS/MS. LC/MS/MS analysis was performed using Vanquish Binary Flex system equipped with TSQ Altis (Thermo Fisher Scientific). The PK parameters of the plasma concentrations of S-217622 were calculated using Phoenix WinNonlin (Ver. 8.1, Certara, L.P.). The plasma concentrations of all dosing groups in the *in vivo* SARS-CoV-2 infection and treatment studies were simulated by non-parametric analysis from plasma concentration data obtained in the PK study. The following PK parameters were calculated based on the non-compartmental method.

#### Metabolic stability studies

Rat liver microsomes (pool of 5, male) were purchased from the Jackson Laboratory Japan, Inc. (Yokohama, Japan) or Charles River Japan, Inc. (Yokohama, Japan). Human liver microsomes (HLM, pool of 15, male and female) were purchased from Sekisui XenoTech (Kansas City, KS). Metabolic stabilities of the test compounds in rat and human liver microsomes were determined at 0.5 μM. The compounds were incubated with 0.5 mg protein/mL in suspension in buffer (50 mM Tris-HCl buffer, pH 7.4, 150 mM KCl, 10 mM MgCl_2_, 1 mM NADPH) at 37°C. Microsomal incubations were initiated by adding 100-fold concentrated solution of the compounds. Incubations were terminated by adding a 2-fold volume of organic solvent (MeCN/MeOH = 1:1) after 0 and 30 min of incubation at 37°C. The precipitation protein was removed by centrifugation. The supernatants were analyzed by liquid chromatography tandem mass spectrometry (LC/MS/MS). LC/MS/MS was performed using LCMS-8060 (SHIMADZU CORPORATION, Kyoto). All incubations were conducted in duplicate, and the percentage of compound remaining at the end of the incubation was determined from the LC/MS/MS peak area ratio.

#### Rat PK studies

The animal study protocol was approved by the director of the institute after reviewing the protocol by the Institutional Animal Care and Use Committee in terms of the 3R (Replacement/Reduction/Refinement) principles.

Rat PK studies were done at Shionogi Pharmaceutical Research Center (Osaka, Japan). Eight-week-old male Sprague-Dawley rats were purchased from Charles River Laboratories. For oral administration, the dosing vehicle was dimethyl sulfoxide/0.5% methylcellulose (400 cP) = 1:4. The compound was orally administered at 1–2 µmol/5 mL/kg (n = 2) under non-fasted conditions. Blood samples (0.2 mL) were collected with 1mL syringes containing anticoagulants (EDTA-2K and heparin) at 0.5, 1, 2, 4, 8, and 24 h after dosing. For intravenous administration, compounds were formulated as solutions in dimethyl sulfoxide/propylene glycol (1:1, v/v) and intravenously administered via the tail vein at 0.5–1.0 µmol/mL/kg (n = 2) under isoflurane anesthesia under non-fasted conditions. Blood samples (0.2 mL) were collected with 1-mL syringes containing anticoagulants (EDTA-2 K and heparin) at 3, 10, 30, 60, 120, 240 and 360 min after dosing. Blood samples were centrifuged to obtain plasma samples, which were transferred to each tube and stored in a freezer until analysis. Plasma concentrations were determined by LC/MS/MS following protein precipitation with MeOH or MeCN. LC/MS/MS analysis was performed using a SCIEX Triple Quad 5500 or SCIEX API5000 or SCIEX Triple Quad 5500 (Sciex, Framingham, MA). PK parameters were calculated via non-compartmental analysis.

#### Dog/Monkey PK studies

PK experiments in dogs and monkeys were conducted in accordance with the guidelines provided by AAALAC. The animal study protocol was approved by the director of the institute after reviewing the protocol by the Institutional Animal Care and Use Committee in terms of the 3R (Replacement/Reduction/Refinement) principles.

Dog PK studies were done at Shionogi Pharmaceutical Research Center (Osaka, Japan), and Monkey PK studies were done at Shionogi Aburahi Research Center (Shiga, Japan). Male beagles were purchased from Marshall BioResources. Female cynomolgus monkeys were purchased from Shin Nippon Biomedical Laboratories, Ltd. or HAMRI CO., Ltd. For oral administration, dosing vehicles were 0.5% methylcellulose (400 cP). The compound was orally administered at 3 mg/2 mL/kg (n = 3) under non-fasted conditions. Blood samples (0.3 mL) were collected with 1-mL syringes containing anticoagulants (EDTA-2K and heparin) at 0.25, 0.5, 1, 2, 4, 8 and 24 h after dosing. For intravenous administration, compounds were formulated as solutions in dimethyl acetamide/ethanol/20% HP-β-CD in carbonate buffer (pH 9.0) (2:3:5, by vol.) and intravenously administered via forelimb or hindlimb vein at 0.1 mg/0.2 mL/kg (n = 2) under non-fasted conditions. Blood samples (0.2 mL) were collected with 1-mL syringes containing anticoagulants (EDTA-2 K and heparin) at 2, 5, 15, 30, 60, 120, 240, 480 and 1440 min after dosing. Blood samples were centrifuged to obtain plasma samples, which were transferred to each tube and stored in a freezer until analysis. Plasma concentrations were determined by LC/MS/MS following protein precipitation with MeOH or MeCN. LC/MS/MS analysis was performed using a SCIEX API5000 or SCIEX Triple Quad 6500 or Triple Quad™ 6500+ (Sciex, Framingham, MA). PK parameters were calculated by non-compartmental analysis.

#### Virtual screening

As a target structure for virtual screening, we retrieved the crystal structure of the SARS-CoV-2 3CL^pro^ in complex with a non-covalent inhibitor, X77 (PDB-ID: 6W63)^19^, from PDB. First, the structure was prepared using Protein Preparation Wizard^28^. Missing atoms and side chains were added, and the ionization states of the amino acids were calculated using Epic^29, 30^. Hydrogen bond networks were optimized, and energy was minimized with a heavy atom restraint of 0.3 Å. All water molecules were removed from the crystal structure, and the docking grid was set to the center of the bound ligand of X77. An in-house compound library was preprocessed by Ligprep^31^ before docking. Virtual screening was performed via Glide^32, 33^ in SP mode. The generated docking poses were filtered by the predefined pharmacophores using Phase^34, 35^. The pharmacophores were set as the acceptor sites with the sidechain NH donor of His163 in the S1 pocket, the lipophilic site in the S2 pocket and the acceptor site with the Glu166 main-chain NH. Finally, the 300 top-scoring compounds that matched all pharmacophores were selected for enzymatic assays. These procedures were conducted using Schrödinger Drug Discovery Suite 2019-4.

#### Expression and purification of SARS-CoV-2 3CL^pro^ protein

The SARS-CoV-2 3CL^pro^ (1-306) containing an *N*-terminal 10-histidine tag followed by a thrombin cleavage site and the SARS-CoV-2 3CL^pro^ (1-306) containing a thrombin cleavage site followed by a C-terminal 10-histidine tag were cloned into pET15b vectors. Two 3CL^pro^ constructs were expressed and purified in the same manner as below. *E. coli* strain BL21 Star (DE3) (Thermo Fisher Scientific) was transformed by the expression plasmid, and then precultivated in LB medium containing 100 mg/mL of ampicillin sodium salt. Six milliliters of preculture was inoculated into 600 mL of fresh TB medium supplemented by 100 mg/mL of ampicillin sodium salt in a 2-L flask with baffles. After vigorous shaking at 37°C, 1 mM IPTG was added for the induction when the optical density (OD)_600_ reached 1.0. After induction for 16 h at 16°C, the cells were harvested by centrifugation.

Cells expressing SARS-CoV-2 3CL^pro^ were resuspended and sonicated. The clarified lysate was subjected to HisTrap FF 5 mL (Cytiva) equilibrated with 20 mM Tris-HCl (pH 8.0), 300 mM NaCl, 1 mM DTT, and 20 mM imidazole, and the proteins were eluted with a linear concentration gradient of imidazole (20–500 mM). Fractions containing SARS-CoV-2 3CL^pro^ were collected and mixed with thrombin His-tag at 4°C overnight to remove the N- or C-terminus. Thrombin-treated SARS-CoV-2 3CL^pro^ was applied to HisTrap FF 5 mL (Cytiva) to remove proteins with uncleaved His-tags. The flow-through fraction was applied to a Superdex 200 16/60 (Cytiva) equilibrated with 20 mM HEPES (pH 7.5), 150 mM NaCl, and 1 mM DTT, and the fraction containing the major peak was collected.

#### Co-crystallization of SARS-CoV-2 3CL^pro^ with compound 1 and 3 (S-217622), diffraction data collection, and structure determination

C-terminal His-tag free SARS-CoV-2 3CL^pro^ protein (4.4 mg/mL) was incubated with 500 mM compound **1** for 1 h at room temperature, and the complexes were crystallized by sitting-drop vapor diffusion at 20°C.

The crystal of the compound **1** complex was grown with buffer containing 0.2 M ammonium citrate tribasic, pH 7.0, with 20% (w/v) PEG 3350.

N-terminal His-tag-free SARS-CoV-2 3CL^pro^ protein (4.6 mg/mL) was incubated with 500 mM of S-217622 for 1 h at room temperature, and the complexes were crystallized by sitting-drop vapor diffusion at 20°C. The S-217622 complex crystal was grown with buffer containing 0.1 M BIS-Tris, pH 6.5, with 2.0 M ammonium sulfate.

X-ray diffraction data were collected using a Rigaku HyPix6000C detector mounted on a Rigaku FR-X rotating anode generator. Data were processed by CrysAlis Pro^36^. The structures were determined by molecular replacement using MOLREP^37^ with the SARS-CoV-2 3CL^pro^-inhibitor complex (PDB-ID 6LU7) as a search model^38^. Iterative model-building cycles were performed with COOT^37^ and refined using REFMAC^39^. The data collection and structure refinement statistics were summarized in Table S5.

## Supporting Information

^1^H-NMR and ^13^C-NMR spectra of synthetic compounds, HPLC traces of compounds **1**-**3**, and S-217622 (**3**) fumaric acid co-crystal. Experimental Procedures for *in vitro* safety, supporting tables (Tables S1-S3) for the *in vitro* antiviral activities of S-217622 (**3**), and tables for the *in vitro* safety experiments (Table S4) and crystallography data collection and refinement statistics (Table S5).

## Accession Code

The coordinates and structural factors of SARS-CoV-2 3CL^pro^ in complex with **1** and **3** (S-217622) have been deposited into PDB with accession numbers 7VTH and 7VU6, respectively. Authors will release the atomic coordinates and experimental data upon article publication.

## Author contributions

Y.U., S.U., and K.N. contributed equally to this paper.

Conceptualization: T.K., Y.T.; Methodology: S.Y., H.N., Y.Y., S.T.; Formal Analysis: S.U., K.N.;H.N.,Y.Y., S.Y., S.K., T.M., T.K., Y.T.; Investigation: Y.U., S.U., K.N., H.N., Y.Y., S.Y., Y.M., Y.T., K.K., T.S., K.K., A.N., S.K., T.S., S.T., K.U., S.A., A.S.; Resources: M.S., Y.O., H.S.; Writing – Original Draft Preparation: Y.U., S.U., K.N., H.N., Y.T.; Writing – Review & Editing: Y.U., S.U., K.N., H.N., T.S., M.S., Y.O., H.S., T.S., T.K., Y.T.; Visualization: Y.U., S.U., H.N., Y.M., S.K., Y.T.; Project Administration: Y.U., S.U., K.N., J.N., Y.Y., S.Y., Y.M., T.M., S.A.,.T.S., T.K., Y.T.; Supervision: T.K., Y.T.

## Notes

SHIONOGI has applied for a patent covering **1**, **2**, and **3** (S-217622). Y.U., S.U., K.N., H.N., Y.Y., S.Y., Y.M., Y.T., K.K., T.S., K.K., A.N., S.K., T.S., S.T., K.U., T.M., S.A., A.S., T.S., T.K., and Y.T. are employees of SHIONOGI & Co., Ltd. S.U., K.N., H.N., Y.M., Y.T., K.K., T.S., K.K., S.K., TS, S.T., K.U., T.S., and T.K. are shareholders in SHIONOGI & Co., Ltd. M.S., Y.O., and H.S. are financially supported by the joint research fund from SHIONOGI & Co., Ltd.

## Acknowledgments

We thank all participants and investigator teams in the clinical study. We acknowledge the National Institute of Infectious Diseases (NIID), Dr. Kouichi Morita (Nagasaki University) and Dr. Bart Haagmans (Erasmus University Medical Center) for providing the SARS-CoV-2 strains, SARS-CoV, and MERS-CoV, respectively. We are also grateful to all our colleagues who participated in the COVID-19 antiviral program at SHIONOGI: Yasushi Hasegawa, Masahiro Masuda, Rina Yasui, Misato Kitamura, Keisuke Mizote, Kotaro Nagatani, Shomitsu Maeno, Tatsuya Tanaka, Azusa Okano, Akinari Sumita, Masayuki Takamatsu, Manabu Kato, Hiroyuki Meichin, Yu Takahashi, Shinya Hisakawa, Yoshihide Sugata, Takao Oyama, Shinichiro Hara, Atsuhiro Iimuro, and Eiichi Kojima, for the compound synthesis; Tetsuya Miyano for the physicochemical studies; Masayoshi Ogawa for structural analysis of the compounds; Akira Kugimiya, Akira Ino, and Kenji Yamawaki for scientific discussion and advice; Takao Shishido, Keita Fukao, Takayuki Kuroda, Masaaki Nakashima, Ryuichi Yano, Yoko Kajiwara, Keiichi Taniguchi, Masaaki Izawa, Shinji Kusakabe, Sachi Takahara, Keiko Baba, and Shigeru Miki for the pharmacological studies; Junji Yamane for X-ray crystallography analysis; Shinpei Yoshida, Yukari Tanaka, Ryoko Oka, and all members of Drug Metabolism & Pharmacokinetics 1&2 group for the DMPK studies; and Yoko Nishimura, Keigo Matsuyama, Sho Hasegawa, Chinami Nekomoto, and Kayoko Kanasaki for the safety studies. We also thank Shionogi Technoadvance Research Co., Ltd. for the compound supplies and technical support in the pharmacological studies. We thank Traci Raley, MS, ELS, from Edanz (https://jp.edanz.com/ac) for editing a draft of this manuscript.

## Abbreviations

CDI: *N*,*N*’-carbonyldiimidazole
DBU: 1,8-diazabicyclo[5.4.0]-7-undecene
DMA: *N*,*N*-dimethylacetamide
DMEM: dulbecco’s modified eagle medium
DMSO: dimethyl sulfoxide
DTT: 1,4-dithiothreitol
ESI: electrospray ionization
EtOAc: ethyl acetate
HATU: 1-[bis(dimethylamino)methylene]-1*H*-1,2,3-triazolo[4,5-*b*]pyridinium 3-oxide hexafluorophosphate
LCMS: liquid chromatography mass spectrometry
LHMDS: lithium bis(trimethylsilyl)amide
HRMS: high resolution mass spectrometry
MeCN: acetonitrile
TFA: trifluoroacetic acid
THF: tetrahydrofuran.

