## Supporting information for "Discovery of S-217622, a Non-Covalent Oral SARS-CoV-2 3CL Protease Inhibitor Clinical Candidate for Treating COVID-19"

**Contents**

- $^{1}$H-NMR and $^{13}$C-NMR spectra for synthesized compounds……………………………………..1
- HPLC traces of **1-3** (S-217622)………………………………………………………...…...13
- Experimental Procedures for *in vitro* safety…………………………………………………...17
- Table S1. The calculated $IC_{50}$ values for S-217622 to SARS-CoV-2 3CL protease activity in each experiment……………………………………………………………………………………..20
- Table S2. $EC_{50}$ values of S-217622 and remdesivir on cytopathic effects in SARS-CoV-2 3CL-infected VeroE6/TMPRSS2 cells…………………………………………………………………...20
- Table S3. $CC_{50}$ values of S-217622 and remdesivir on cytopathic effects in VeroE6/TMPRSS2 cells………………………………………………………………………………………………21
- Table S4. *In vitro* safety profiles of S-217622……………………………………………………21
- Table S5. Diffraction data and refinement statistics for SARS-CoV-2 $3CL^{pro}$ complexed with compound **1** and compound **3** (**S-217622**)………………………………………………………..22

### $^1$H-NMR and $^{13}$C-NMR spectra for synthesized compounds

$^1$H NMR spectra of **5** in DMSO-$d_6$

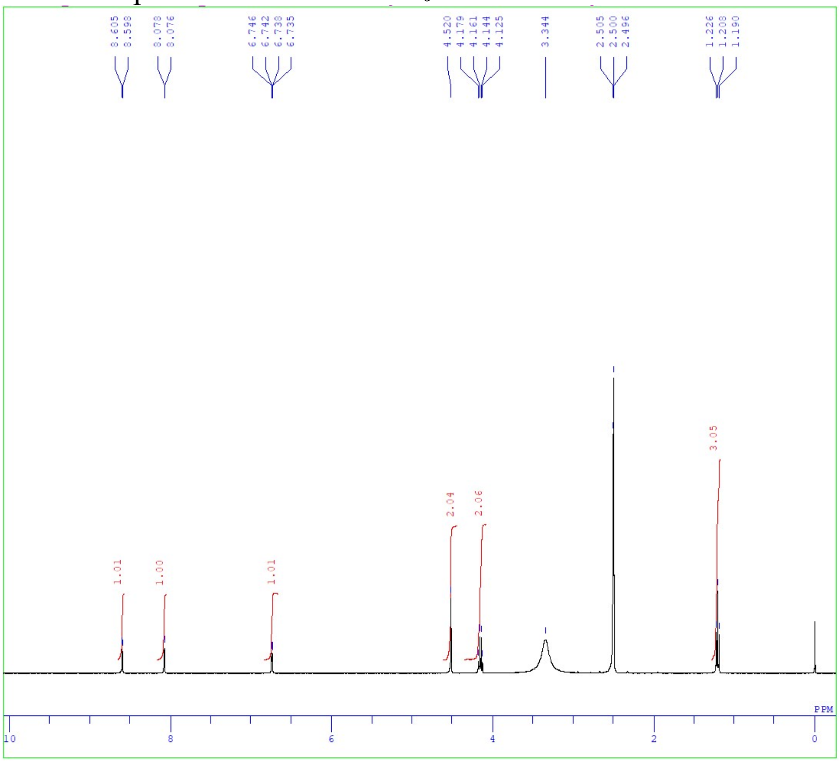


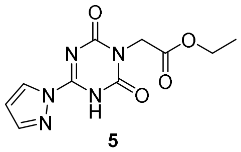

**5**

$^{13}$C NMR spectra of **5** in DMSO-$d_6$

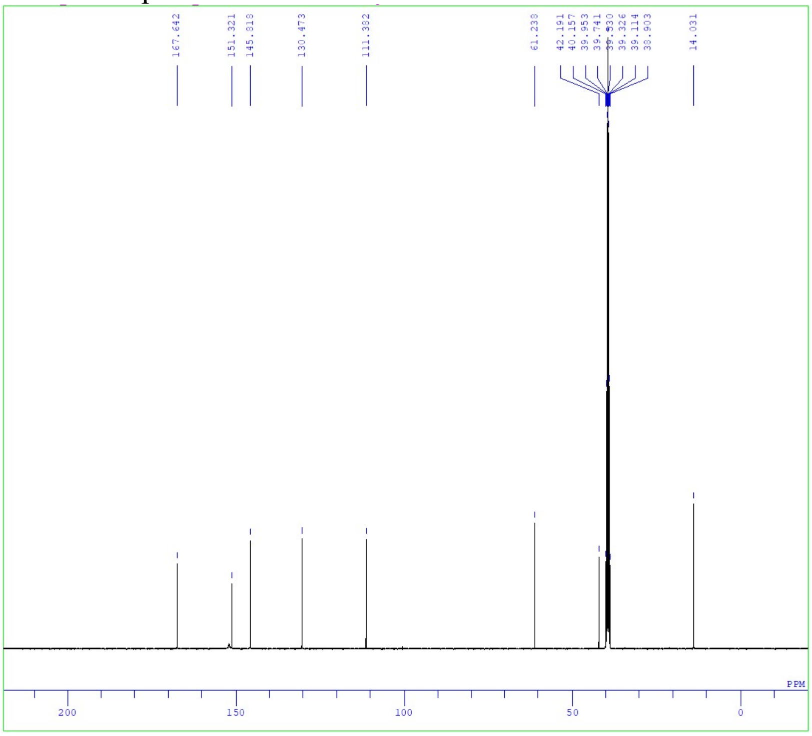


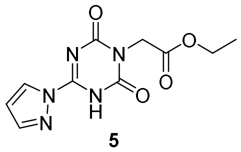

**5**

[1]H NMR spectra of **7** in $CDCl_3$

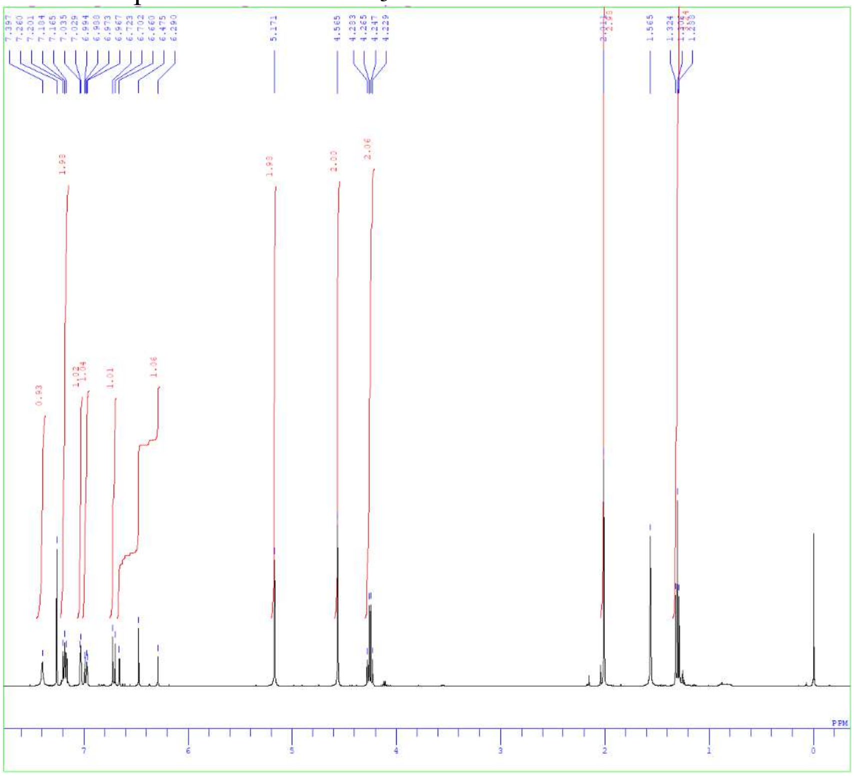

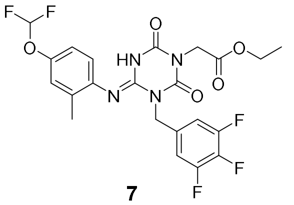

**7**

[13]C NMR spectra of **7** in $CDCl_3$

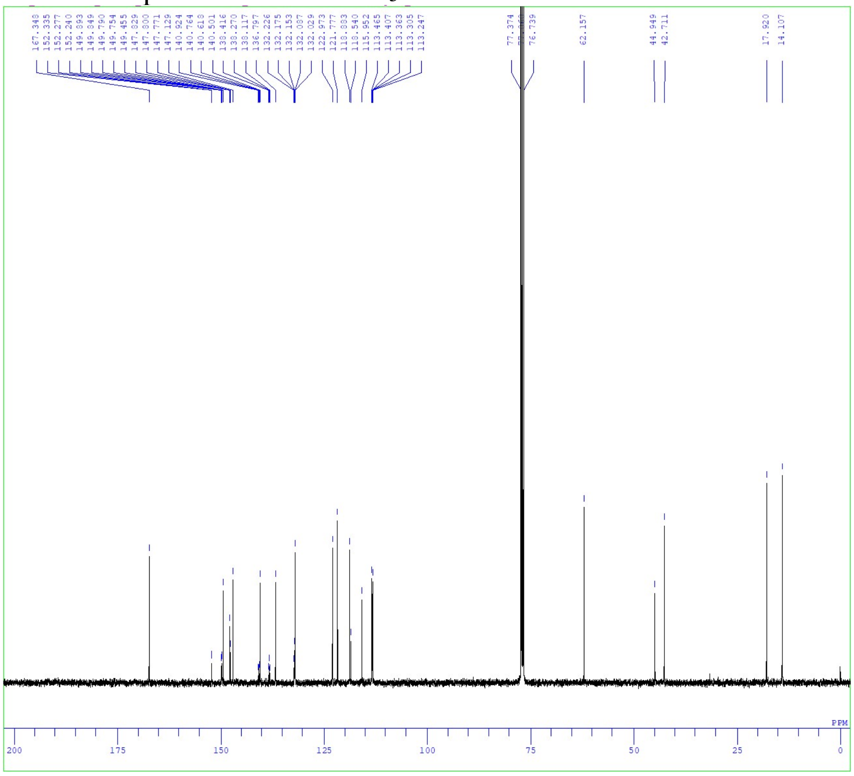

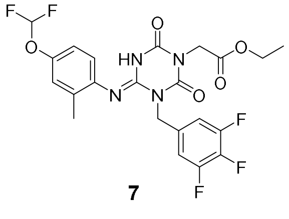

**7**

$^{1}$H NMR spectra of **1** in $CDCl_3$

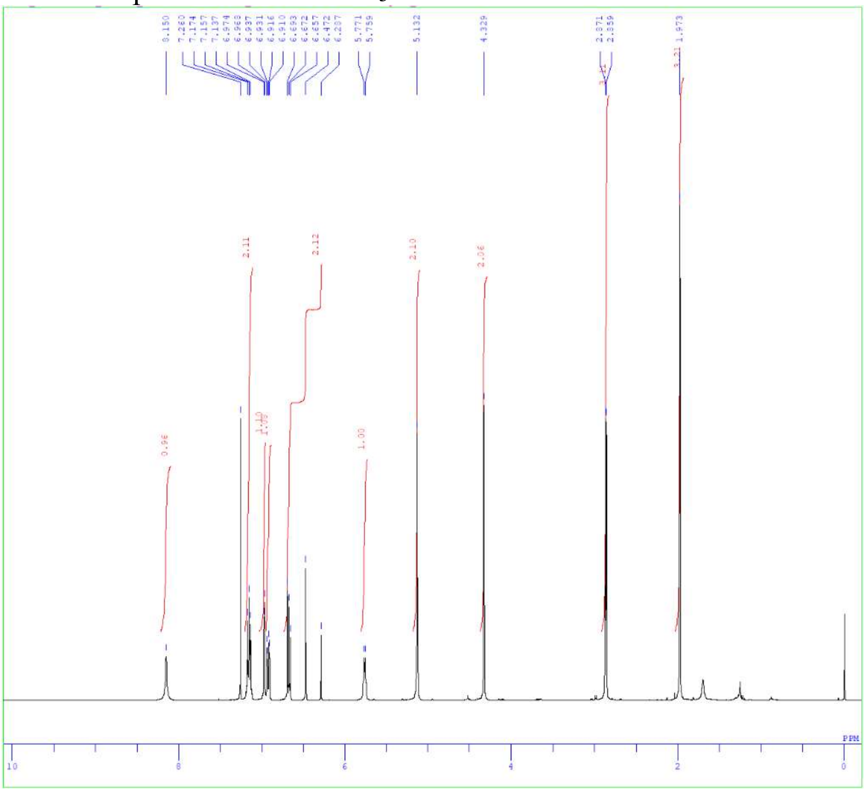

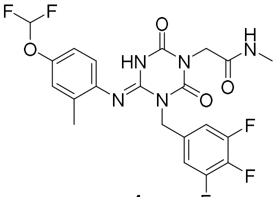

**1**

$^{13}$C NMR spectra of **1** in $CDCl_3$

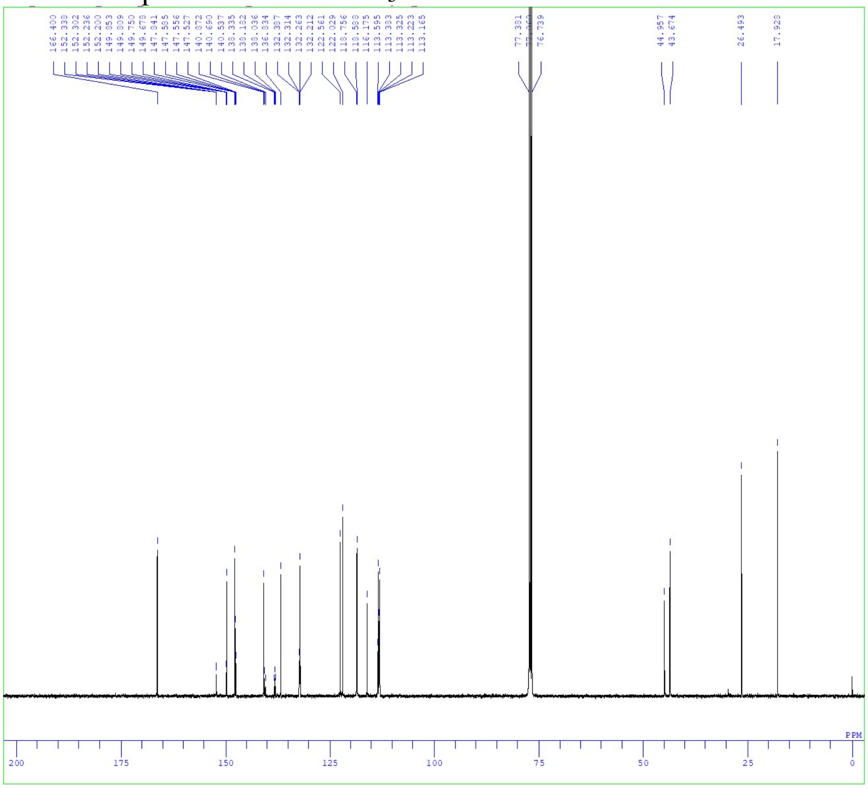

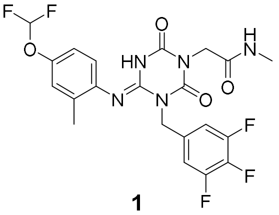

**1**

[1]H NMR spectra of **8** in DMSO-$d_6$ with DCl in $D_2O$

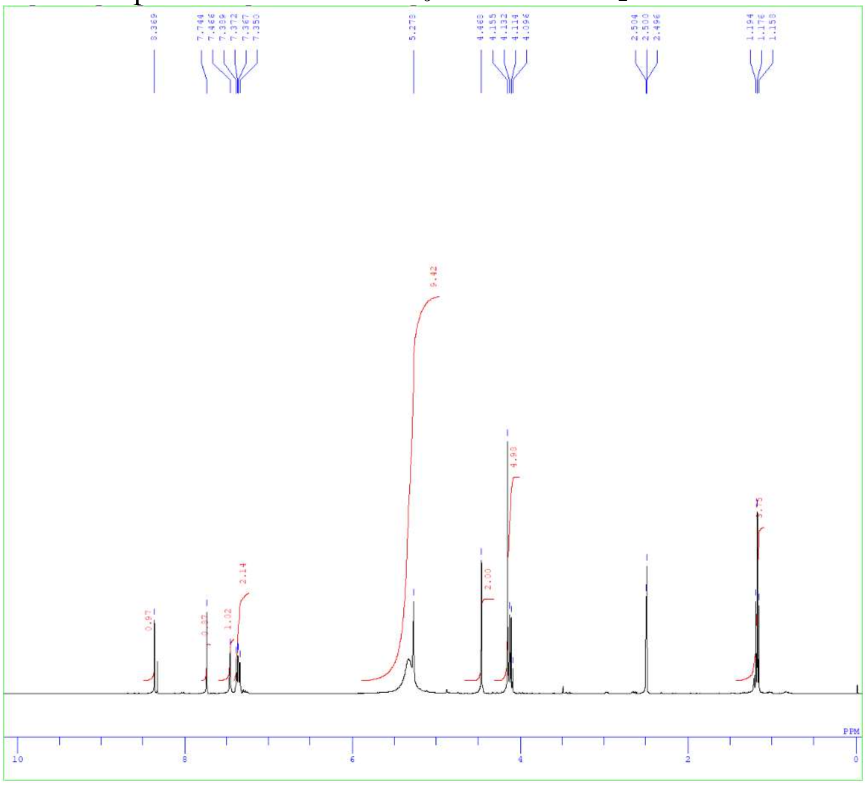

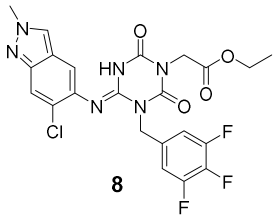

**8**

[13]C NMR spectra of **8** in DMSO-$d_6$ with DCl in $D_2O$

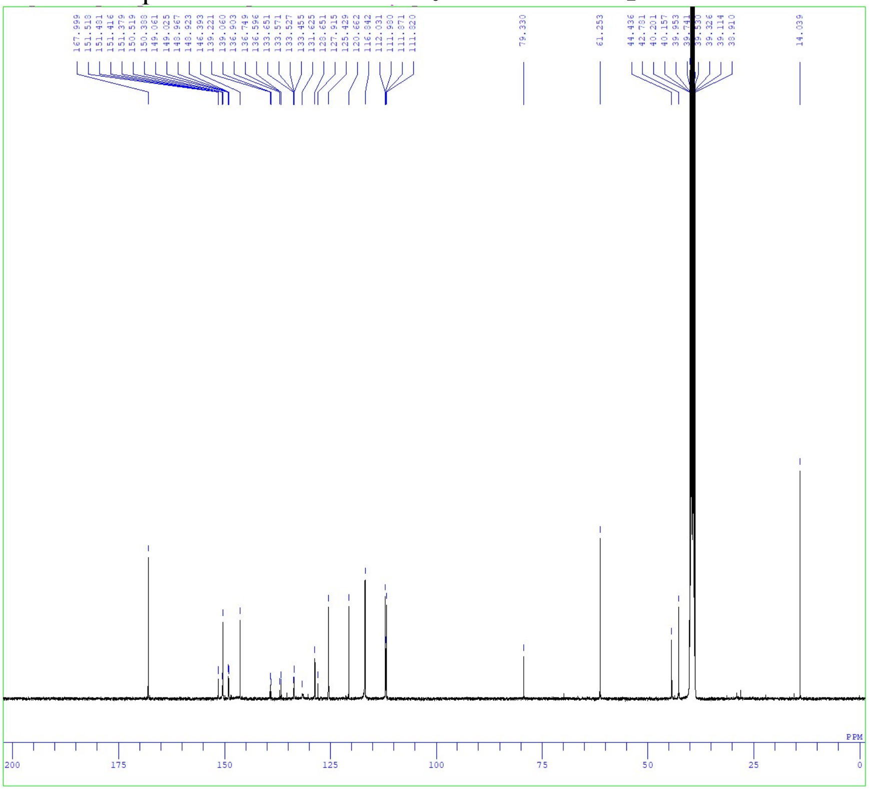

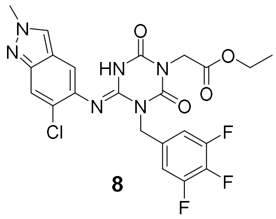

**8**

[1]H NMR spectra of **2** in DMSO-$d_6$ with DCl in $D_2O$

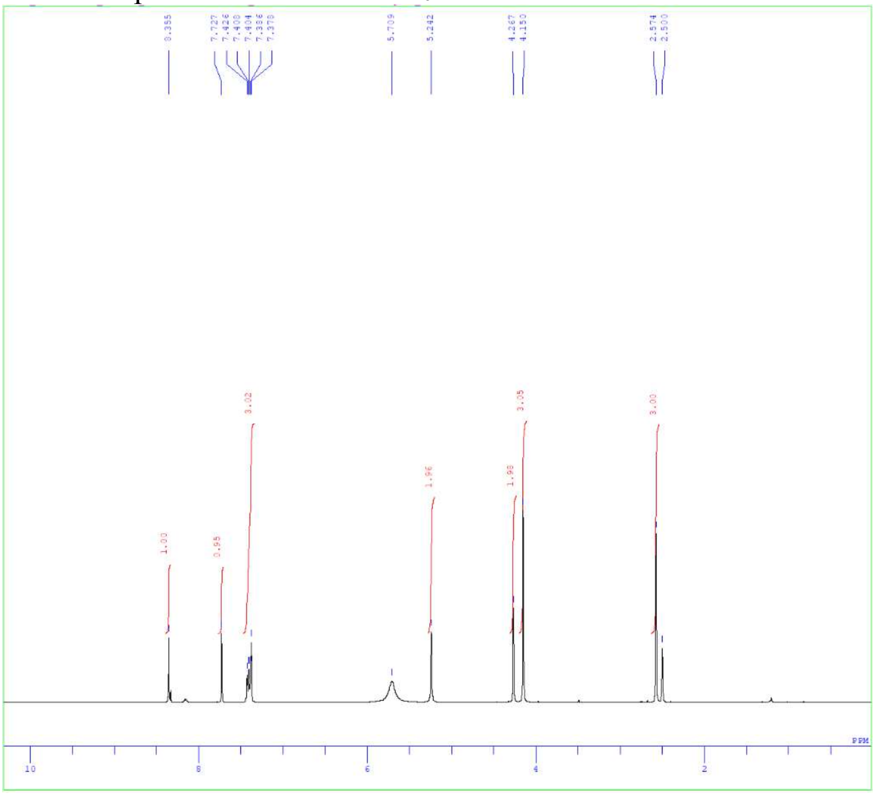

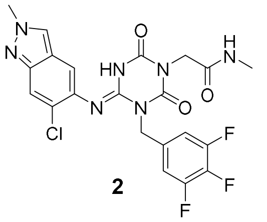

**2**

[13]C NMR spectra of **2** in DMSO-$d_6$ with DCl in $D_2O$

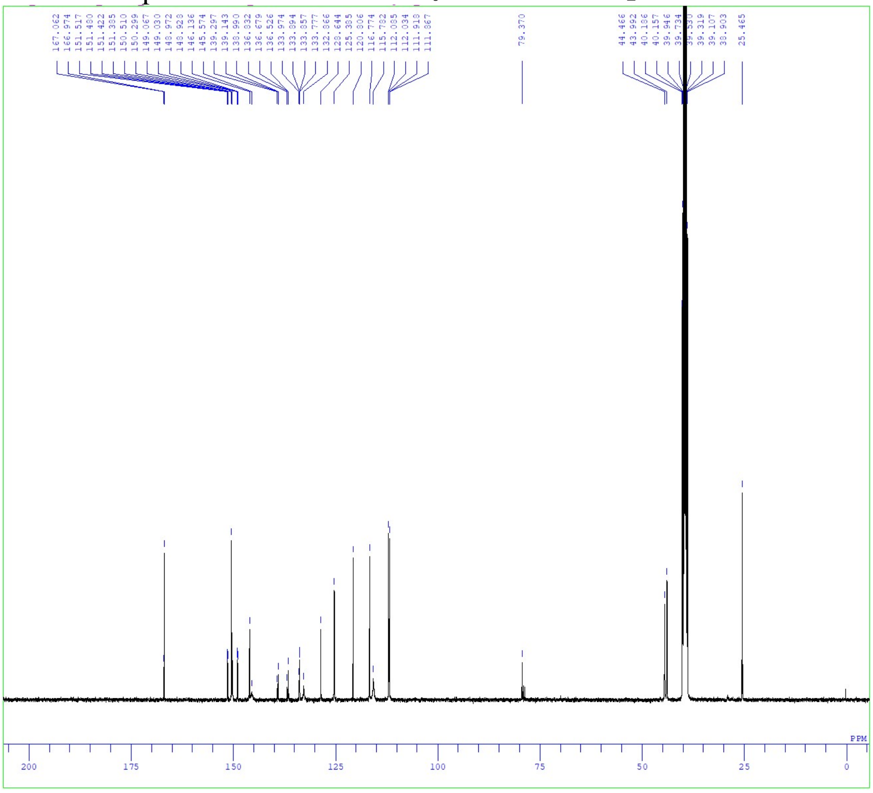

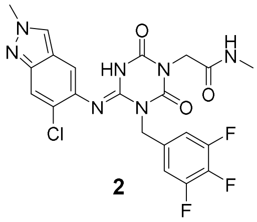

**2**

$^{1}$H NMR spectra of **9** in $CDCl_3$

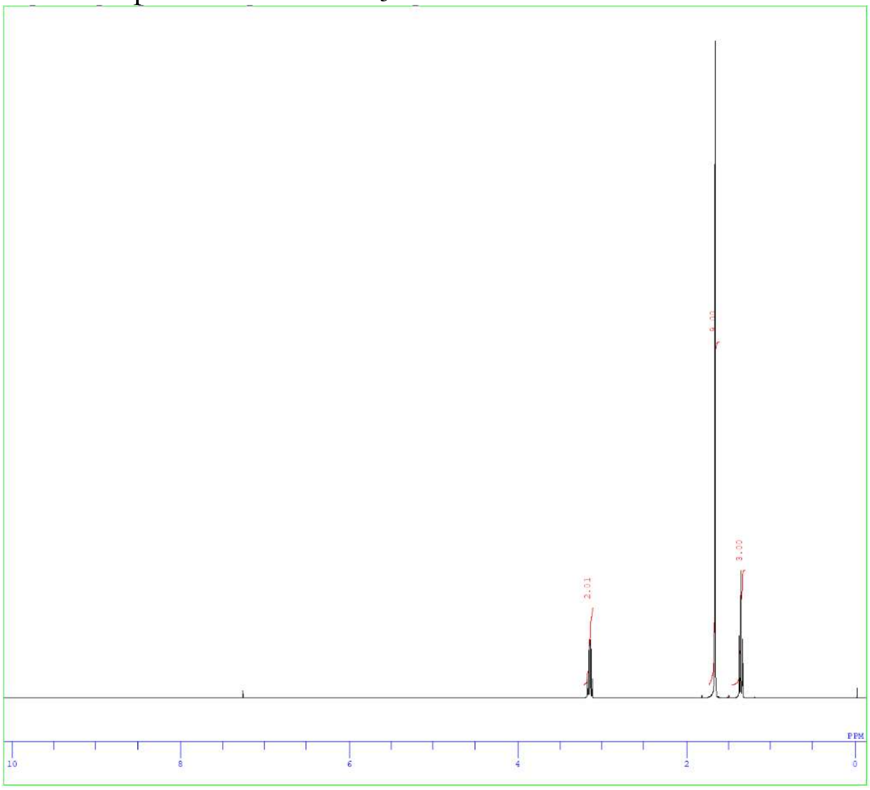


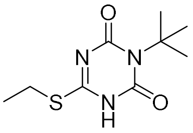


**9**

$^{13}$C NMR spectra of **9** in $CDCl_3$

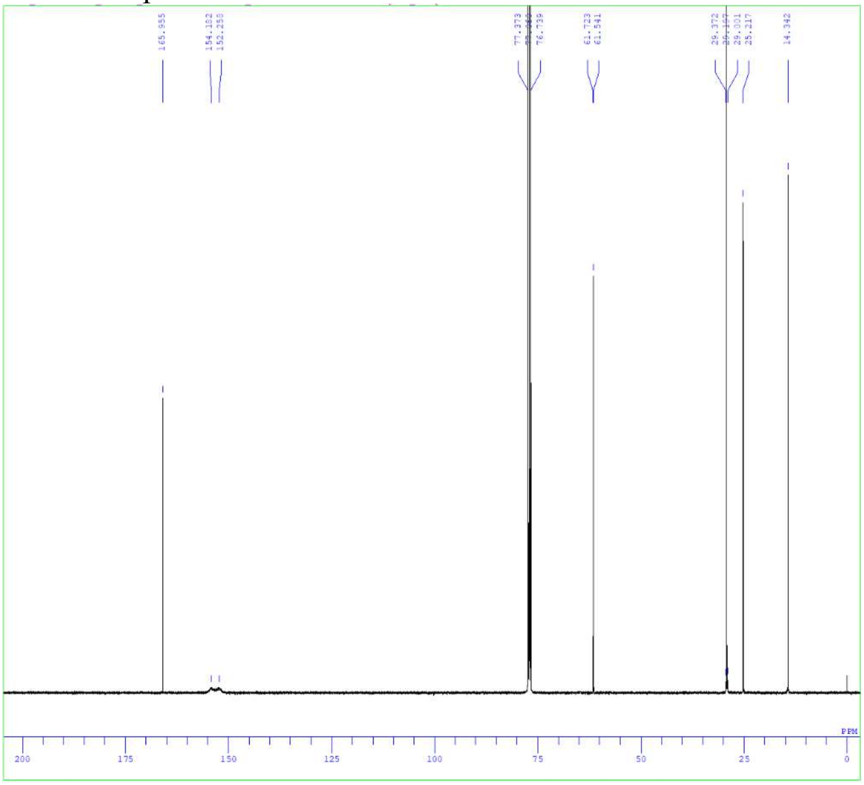


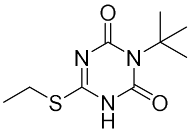


**9**

$^{1}H$ NMR spectra of **10** in $CDCl_3$

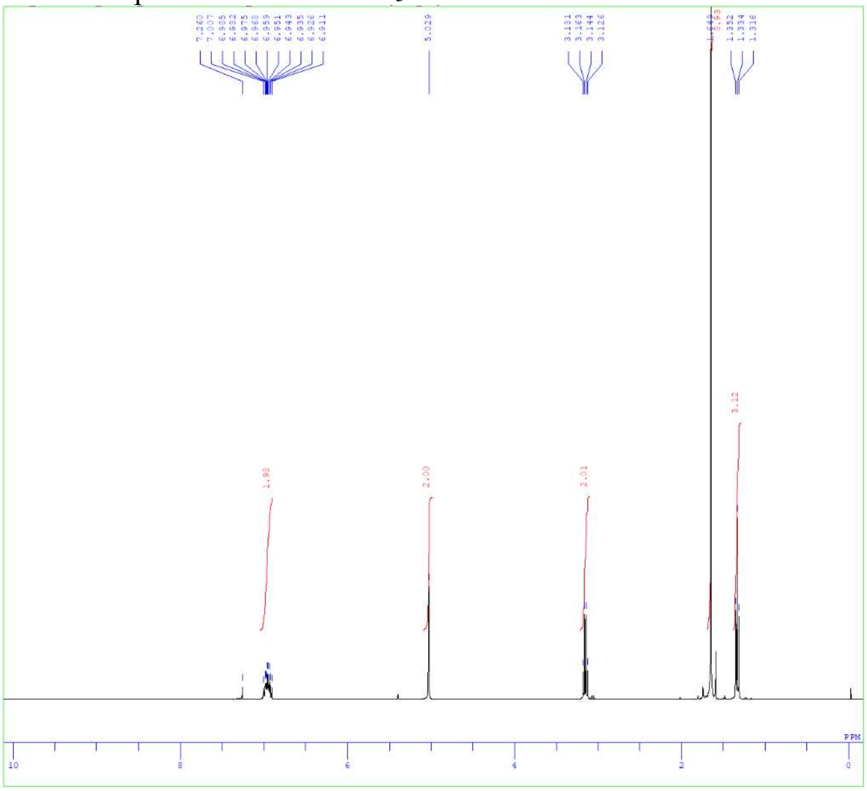

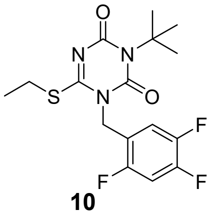


$^{13}C$ NMR spectra of **10** in $CDCl_3$

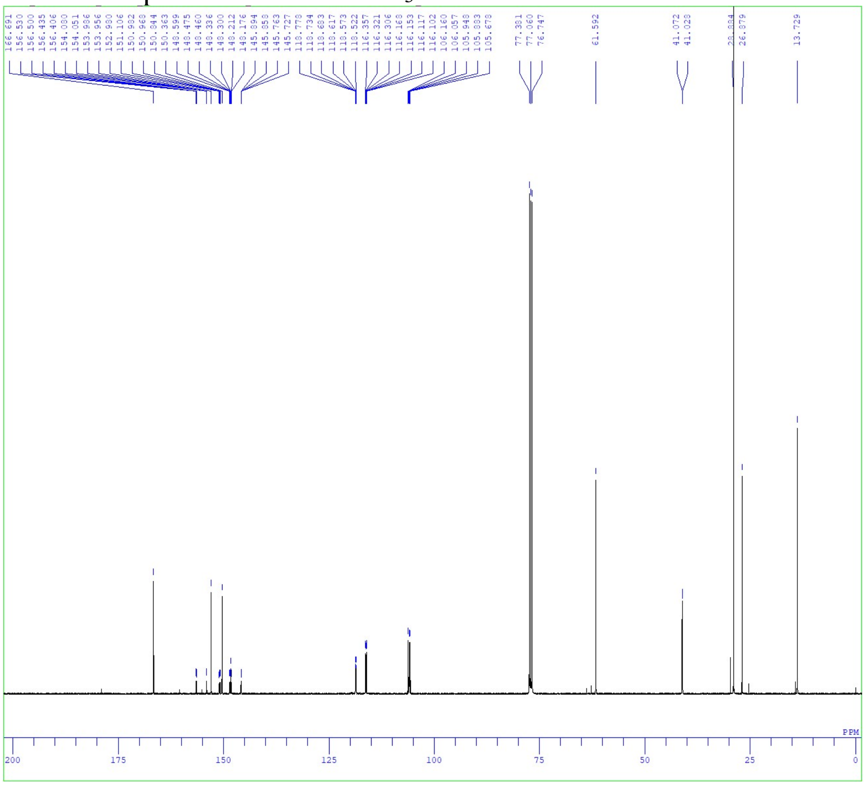

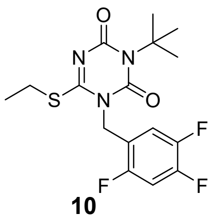

[1]H NMR spectra of **11** in $CDCl_3$

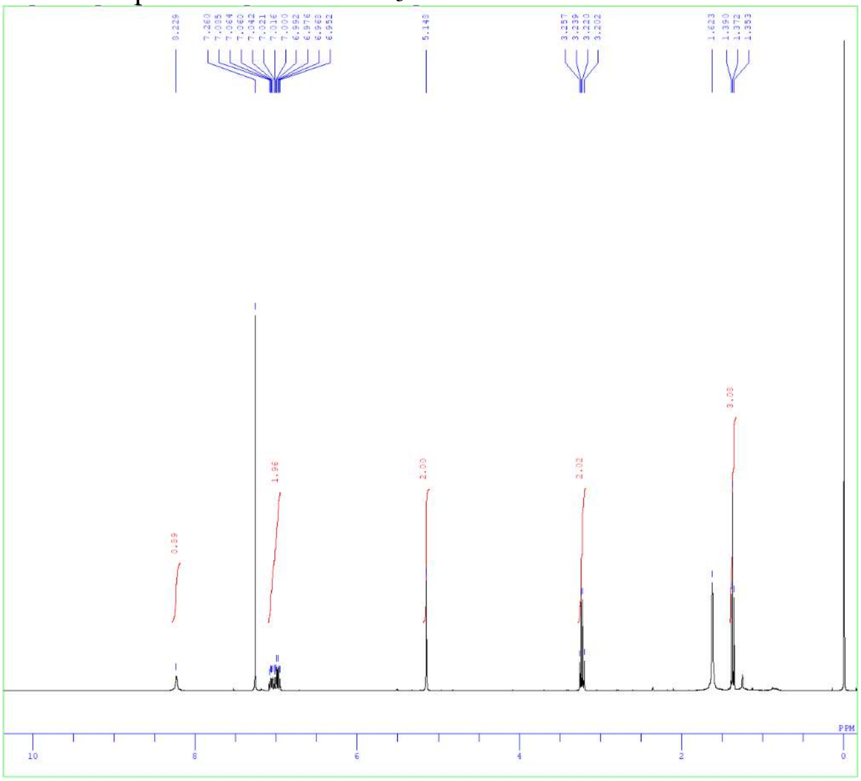

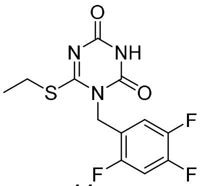

**11**

[13]C NMR spectra of **11** in DMSO-$d_6$

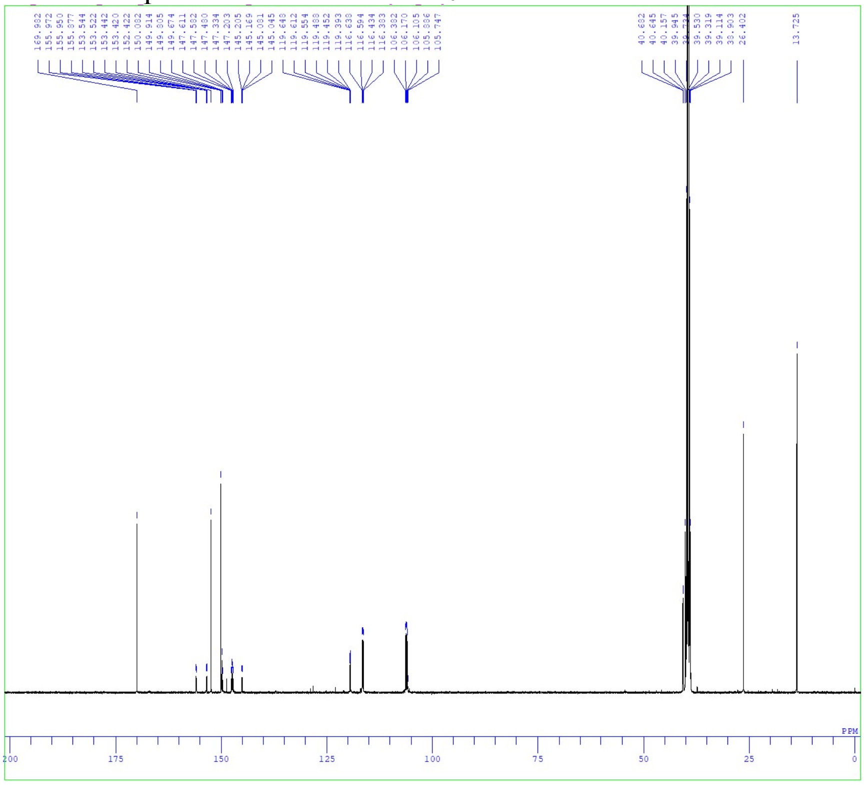

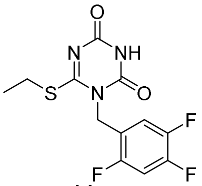

**11**

$^{1}H$ NMR spectra of **12** in $CDCl_3$

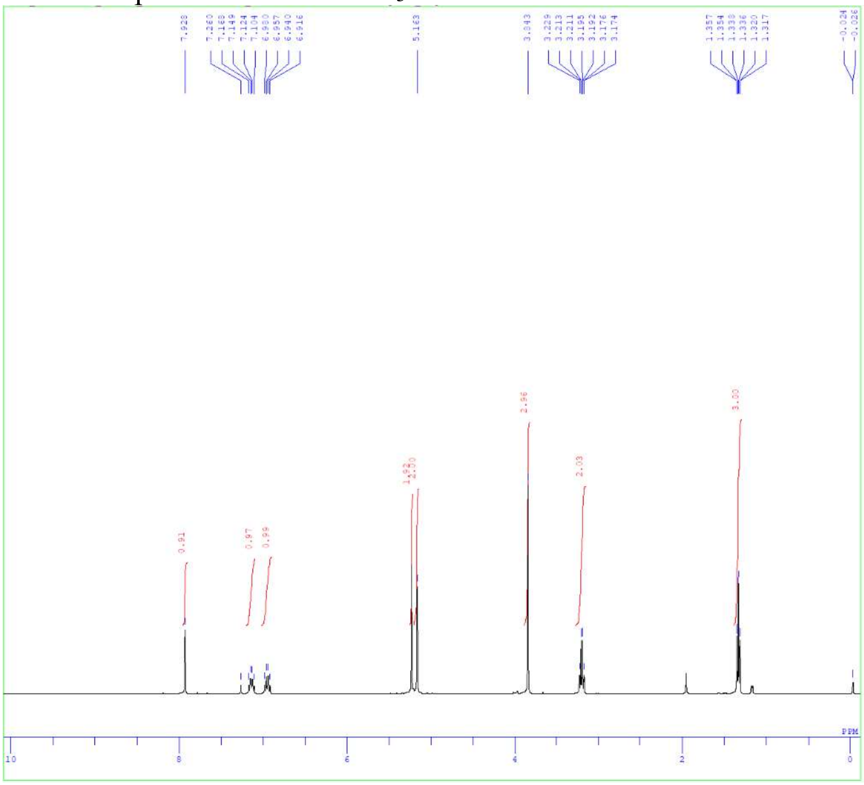

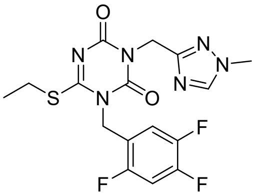

**12**

$^{13}C$ NMR spectra of **12** in $CDCl_3$

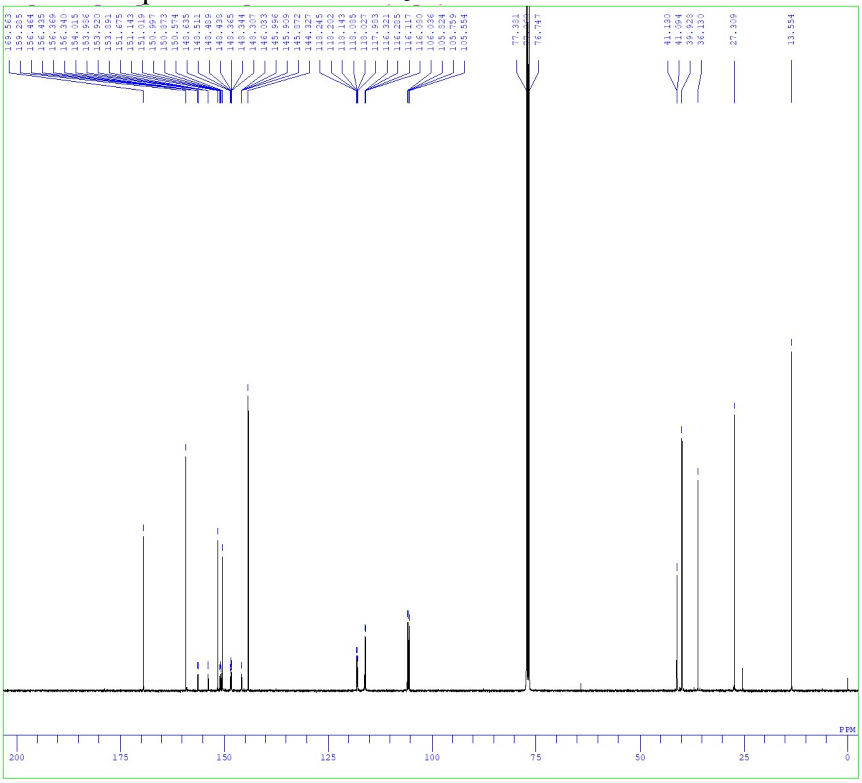

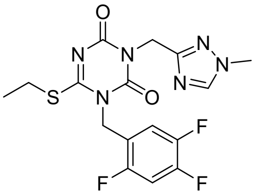

**12**

[1]H NMR spectra of **3** in DMSO-$d_6$ with DCl in $D_2O$

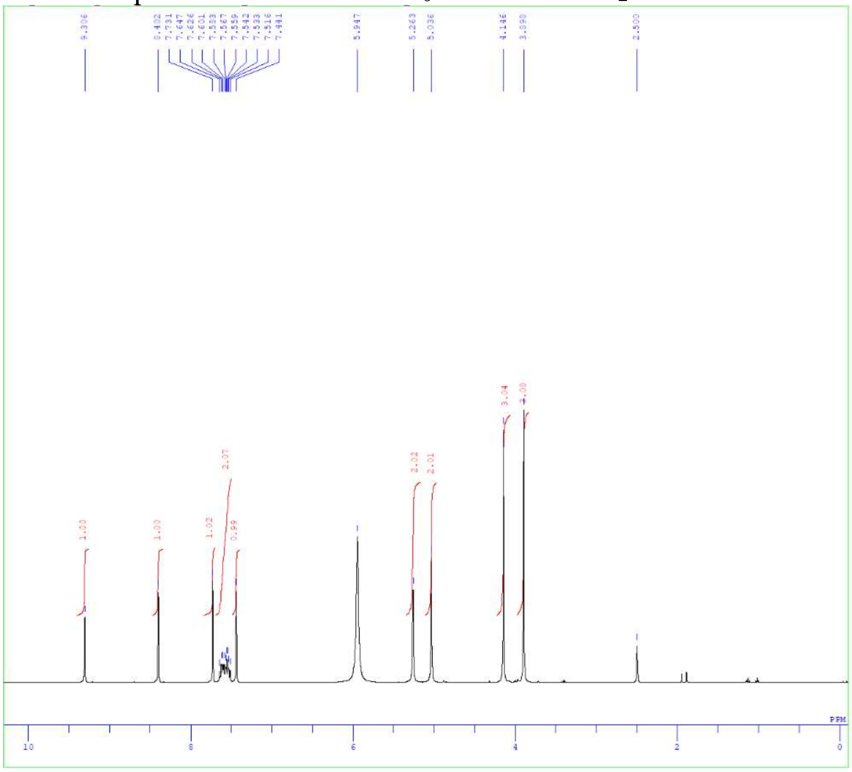


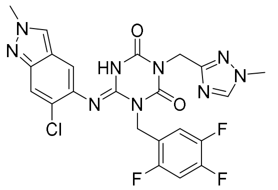


**3**

[13]C NMR spectra of **3** in DMSO-$d_6$ with DCl in $D_2O$

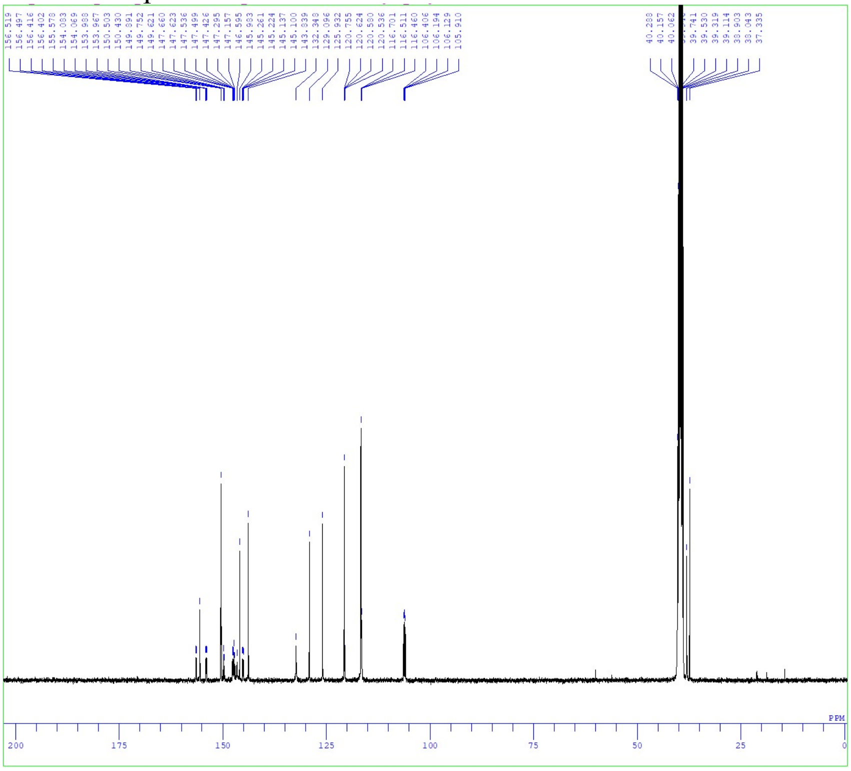


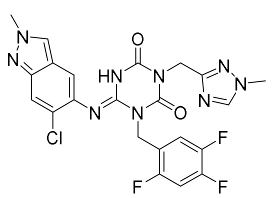


**3**

$^1$H NMR spectra of compound **3 (S-217622)** fumaric acid co-crystal in pyridine-$d_5$

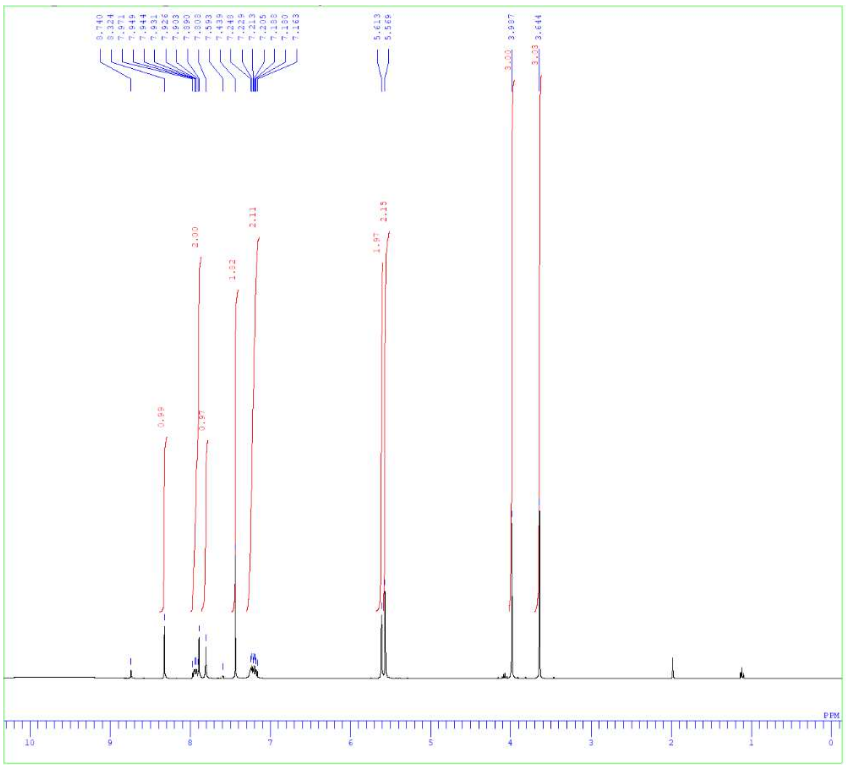

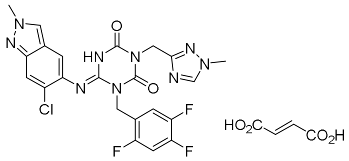

**3 (fumaric acid co-crystal)**

**HPLC traces of compound 1-3**

**Procedure**

Analytical liquid chromatography/mass spectroscopy (LC/MS) was performed on a Shimadzu Shim-pack XR-ODS (C18, 2.2 µm, 3.0 × 50 mm, linear gradient from 10% to 100% B over 3 min, then 100% B for 1 min [A = water + 0.1% formic acid, B = MeCN + 0.1% formic acid], flow rate: 1.6 mL/min) using a Shimadzu UFLC system equipped with a LCMS-2020 mass spectrometer, LC-20AD binary gradient module, SPD-M20A photodiode array detector (detection at 254 nm), and SIL-20AC sample manager.

Compound **1**

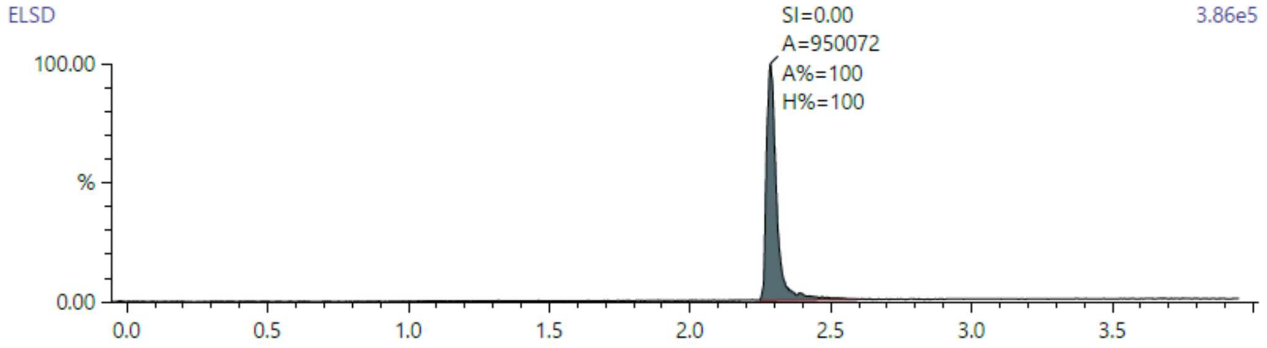


| Pk# | RT | Area | Area % | Height | Height % |
|---|---|---|---|---|---|
| 6 | 2.286 | 950,072 | 100.000 | 376,089 | 100.000 |
| **Totals:** | | **950,072** | **100.000** | **376,089** | **100.000** |

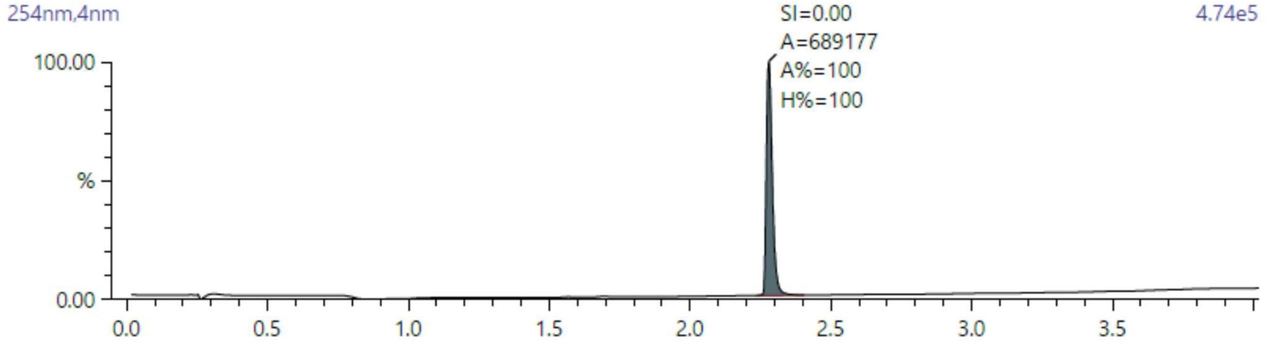


| Pk# | RT | Area | Area % | Height | Height % |
|---|---|---|---|---|---|
| 6 | 2.279 | 689,177 | 100.000 | 474,219 | 100.000 |
| **Totals:** | | **689,177** | **100.000** | **474,219** | **100.000** |

Compound **2**

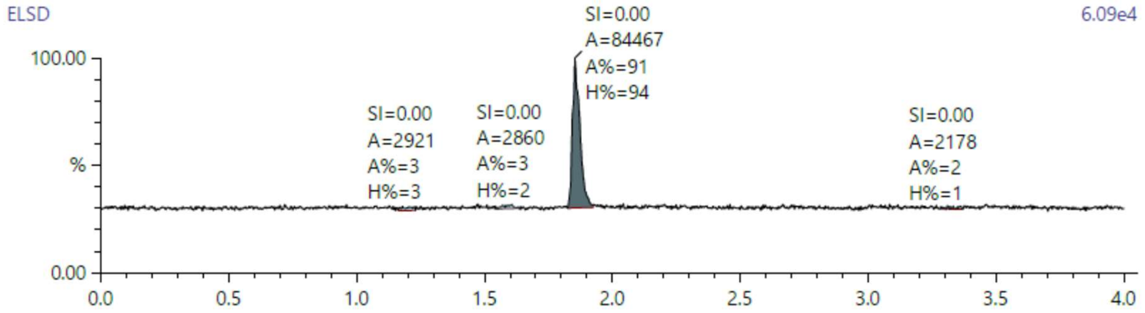


| Pk# | RT | Area | Area % | Height | Height % |
|---|---|---|---|---|---|
| 1 | 1.180 | 2,921 | 3.161 | 1,204 | 2.665 |
| 2 | 1.607 | 2,860 | 3.094 | 1,126 | 2.492 |
| 3 | 1.854 | 84,467 | 91.389 | 42,478 | 94.042 |
| 4 | 3.297 | 2,178 | 2.356 | 362 | 0.801 |
| **Totals:** | | **92,426** | **100.000** | **45,169** | **100.000** |

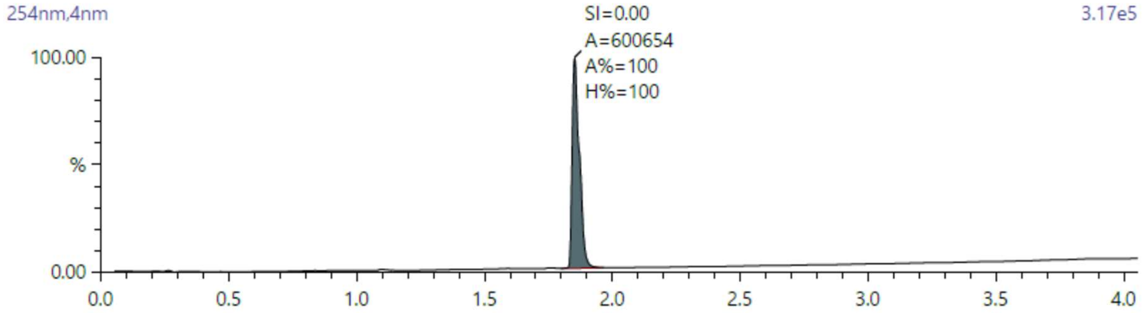


| Pk# | RT | Area | Area % | Height | Height % |
|---|---|---|---|---|---|
| 3 | 1.853 | 600,654 | 100.000 | 312,342 | 100.000 |
| **Totals:** | | **600,654** | **100.000** | **312,342** | **100.000** |

Compound **3** (free form)

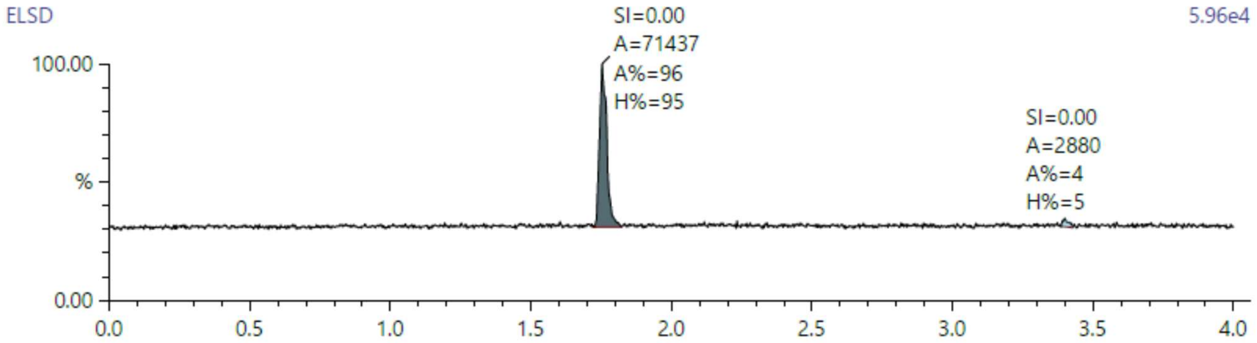


| Pk# | RT | Area | Area % | Height | Height % |
|---|---|---|---|---|---|
| 1 | 1.754 | 71,437 | 96.124 | 41,034 | 95.111 |
| 2 | 3.399 | 2,880 | 3.876 | 2,109 | 4.889 |
| **Totals:** | | **74,317** | **100.000** | **43,144** | **100.000** |

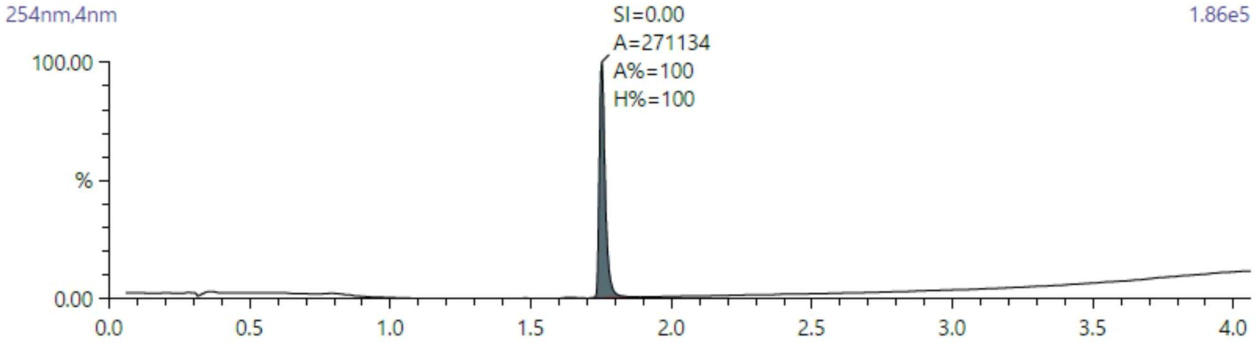


| Pk# | RT | Area | Area % | Height | Height % |
|---|---|---|---|---|---|
| 1 | 1.752 | 271,134 | 100.000 | 189,771 | 100.000 |
| **Totals:** | | **271,134** | **100.000** | **189,771** | **100.000** |

### HPLC chromatogram of 3 (S-217622) fumaric acid co-crystal

Analytical liquid chromatography of S-217622 fumaric acid co-crystal was performed on a ACQUITY UPLC BEH C18 Column (1.7 µm, 2.1 mm I.D.x100 mm, gradient from 10% to 100% B [A = 10 mmol/L $HCOONH_4$ /$H_2O$, B = MeCN], flow rate: 0.3 mL/min) using a Shimadzu Nexera system equipped with LC-20AD binary gradient module and SPD-20AV detector (detection at 255 nm).

Time-program for gradient elution

| Time (min) | Mobile Phase B (%) |
|---|---|
| 0 | 10 |
| 3 | 30 |
| 20 | 30 |
| 26 | 90 |
| 32 | 90 |
| 32.01 | 10 |
| 42 | Stop |

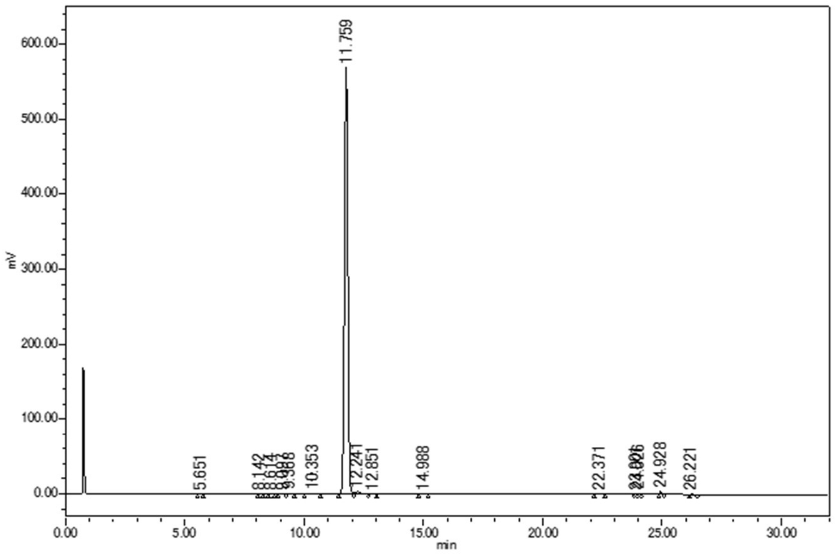


| Retention time (min) | Peak area (µV·sec) | Peak height (µV) | % area |
|---|---|---|---|
| 5.7 | 1179 | 320 | 0.02 |
| 8.1 | 1119 | 212 | 0.02 |
| 8.6 | 1316 | 219 | 0.02 |
| 9.1 | 1095 | 132 | 0.02 |
| 9.4 | 6948 | 987 | 0.12 |
| 10.4 | 14968 | 881 | 0.26 |
| 11.8 | 5559146 | 569193 | 98.21 |
| 12.2 | 45115 | 3668 | 0.80 |
| 12.9 | 1613 | 151 | 0.03 |
| 15.0 | 1242 | 102 | 0.02 |
| 22.4 | 4185 | 364 | 0.07 |
| 23.9 | 1721 | 436 | 0.03 |
| 24.0 | 4907 | 1408 | 0.09 |
| 24.9 | 13517 | 4116 | 0.24 |
| 26.2 | 2513 | 410 | 0.04 |

**Experimental Procedures for *in vitro* safety.**

**Human ether-a-go-go-related gene inhibition assay**

To evaluate an electrocardiogram QT interval prolongation of S-217622, effects on delayed rectifier K+ current (IKr) were evaluated using Chinese hamster ovary (CHO) cells expressing the human ether-a-go-go-related gene channel. The study was conducted in compliance with good laboratory practice regulations.

CHO cells were retained at a membrane potential of −80 mV with a whole-cell patch-clamp system (EPC-10 amplifier/PatchMaster v2.8 software, HEKA Co., Ltd.), and IKr was elicited via repolarization pulse at −40 mV for 2 sec after a depolarization pulse at +20 mV for 1 sec. S-217622 (fumaric acid co-crystal) was dissolved in DMSO and diluted 200-fold with external solution to prepare an objective concentration. The S-217622 concentration levels were 10, 30 and 100 μM. E-4031 at 0.1 μM and the vehicle (0.5% DMSO) were applied as positive and negative controls, respectively.

From the recorded IKr, an absolute value of the tail peak current was measured based on the current value at the resting membrane potential using the whole-cell patch-clamp method. The percentages of the preapplication values in the test substance and control groups were compensated by the mean value of the percentage of the preapplication value in the negative-control group, and the compensated suppression rates were calculated.

Patch-clamp solutions were as follows. Internal solution: KCL: 130mmol/L, $MgCl_2$: 1 mmol/L, MgATP: 5 mmol/L, EGTA: 5 mmol/L, HEPES (4-(2-hydroxyethyl)-1-piperazineethanesulfonic acid): 10 mmol/L, pH=7.2. External solution: NaCl: 137 mmol/L, KCl: 4 mmol/L, $CaCl_2$: 1.8 mmol/L, $MgCl_2$: 1 mmol/L, glucose: 10 mmol/L, HEPES: 10 mmol/L, pH=7.4.

***In vitro* micronucleus test.** To evaluate the clastogenic potential of S-217622, an *in vitro* micronucleus test was conducted using TK6 cells (human lymphoblast-derived) in a short-term treatment with and without a metabolic activation system (S9 mix) and continuous treatment without S9 mix. S9 mix containing 9,000 g of liver supernatant fraction was prepared from Sprague-Dawley rats treated with phenobarbital and 5,6-benzofravone. The study was conducted in compliance with good laboratory practice regulations.

A TK6 cell suspension was used in the absence of metabolic activation in the 3-h and 24-h treatment groups or in the presence of metabolic activation in the 3-h treatment group. Suspensions were mixed with S-217622 (fumaric acid co-crystal) in DMSO solution with S9 mix in the presence of metabolic activation in the 4-h treatment group, then incubated at 37°C. The negative-control (DMSO) and positive-control (mitomycin C, cyclophosphamide monohydrate, or colchicine) substances were prepared concurrently. After the end of the short-term treatment, cells were washed and then incubated in fresh culture medium for 21 h. After incubation, the cells were counted to evaluate cytotoxicity, and the *in vitro* micronucleus test was conducted with S-217622 (fumaric acid co-crystal) at doses of 150–250 μg/mL for short-term treatment and 75–125 μg/mL for continuous treatment based on cytotoxicity. The nucleic acid was stained with acridine orange, and the micronuclear frequency of the specimens was observed under a fluorescence microscope. The test was considered positive when a significant and dose-dependent increase was noted in the number of cells with micronuclei in the test-substance groups compared with the negative-control group under any treatment condition.

**Ames test.** To evaluate the mutagenic potential of S-217622, a bacterial reverse-mutation test was conducted via the preincubation method using five bacterial strains, including *Salmonella typhimurium* (TA98, TA100, TA1535, and TA1537) and *Escherichia coli* (WP2*uvrA*) in the presence or absence of a metabolic activation system (S9 mix). S9 mix containing 9,000 g of liver supernatant fraction was prepared from Sprague-Dawley rats treated with phenobarbital and 5,6-benzofravone. The study was conducted in compliance with good laboratory practice regulations.

S-217622 (fumaric acid co-crystal, DMSO solution) was mixed with S9 mix in the presence of metabolic activation or phosphate buffer in the absence of metabolic activation and 0.1 mL of test strain suspension ($1 \times 10^9$ cells/mL or greater) and incubated at 37°C for 20 min. Then, the mixture with a layer of soft agar containing histidine and biotin or tryptophan was overlaid on minimal glucose agar plates. The negative-control (DMSO) and positive-control (4-nitroquinoline 1-oxide, sodium azide, 9-aminoacridine hydrochloride monohydrate, or 2-aminoanthracene) substances were prepared concurrently. The mutation test was conducted with S-217622 (fumaric acid co-crystal) at 156–5000 μg/plate in TA98, TA100, TA1535, and WP2*uvrA* and at 39.1–5000 μg/plate in TA1537. After incubation at 37°C for 48 h, the revertant colonies were counted and evaluated by comparing them with the negative-control group. The test was considered positive when the number of revertant colonies was concentration-dependently increased and twofold or greater increased over the number of colonies of the negative-control group.

***In vitro* 3T3.** To evaluate the phototoxicity potential of S-217622, an *in vitro* phototoxicity study was conducted with cultured mammalian cells. The study was conducted in compliance with good laboratory practice regulations. A fibroblastic cell line derived from BALB/c mice (BALB/3T3 cells) was cultured in 96-well plates and treated with S-217622 (fumaric acid co-crystal) for 1 h followed by UV-A (5 $J/cm^2$) and UV-B (68.7 $mJ/cm^2$) irradiation. For the comparator, no irradiation was conducted. The phototoxicity test was performed at 0.781–100 μg/mL. A vehicle (DMSO)-treated group and a chlorpromazine hydrochloride-treated group were set as the negative and positive controls, respectively. Cell viability was determined by neutral-red extraction from cells (measurement of absorbance at 540 nm). When the $IC_{50}$ could be determined for both the irradiated and nonirradiated plates, the result was determined from the PIF. When the $IC_{50}$ could not be determined for either the irradiated or nonirradiated plates, the result was determined from the MPE. The judgment criteria are shown below.

No phototoxicity: PIF $< 5$ or MPE $< 0.15$
Phototoxicity: $5 \leq$ PIF or $0.15 \leq$ MPE

**Table S1.** The calculated $IC_{50}$ values for S-217622 (fumaric acid co-crystal) to SARS-CoV-2 3CL protease activity in each experiment.

| **Experiment** | **$IC_{50}$ (μM)** |
|---|---|
| Exp. No.1 | 0.0122 |
| Exp. No.2 | 0.0130 |
| Exp. No.3 | 0.0143 |
| Mean | 0.0132 |
| Standard deviation | 0.0011 |

**Table S2.** $EC_{50}$ values of S-217622 (fumaric acid co-crystal) and remdesivir on cytopathic effects in SARS-CoV-2 3CL-infected VeroE6/TMPRSS2 cells. [a]standard deviation. The mean and SD were calculated from three independent experiments.

| **Strains** | **Pango Lineage** | **$EC_{50}$ (μM)** | | | |
|---|---|---|---|---|---|
| | | **S-217622** | | **Remdesivir** | |
| | | **Mean** | **SD[a]** | **Mean** | **SD[a]** |
| hCoV-19/Japan/TY/WK-521/2020 | A | 0.37 | 0.060 | 1.9 | 0.14 |
| hCoV-19/Japan/QK002/2020 | B.1.1.7 | 0.33 | 0.050 | 0.87 | 0.027 |
| hCoV-19/Japan/QHN001/2020 | B.1.1.7 | 0.31 | 0.070 | 0.97 | 0.14 |
| hCoV-19/Japan/QHN002/2020 | B.1.1.7 | 0.46 | 0.044 | 0.99 | 0.18 |
| hCoV-19/Japan/TY7-501/2021 | P.1 | 0.50 | 0.048 | 2.1 | 0.39 |
| hCoV-19/Japan/TY7-503/2021 | P.1 | 0.43 | 0.00085 | 1.0 | 0.16 |
| hCoV-19/Japan/TY8-612/2021 | B.1.351 | 0.40 | 0.048 | 1.2 | 0.30 |
| hCoV-19/Japan/TY11-927-P1/2021 | B.1.617.2 | 0.41 | 0.014 | 1.6 | 0.22 |
| hCoV-19/Japan/TY38-873/2021 | B.1.1.529 | 0.29 | 0.054 | 1.1 | 0.28 |

**Table S3.** $CC_{50}$ values of S-217622 (fumaric acid co-crystal) and remdesivir against VeroE6/TMPRSS2 cells. [a]standard deviation. The mean and SD were calculated from three independent experiments.

| Substance | $CC_{50}$ (μM) | | | | |
|---|---|---|---|---|---|
| | Exp.1 | Exp.2 | Exp.3 | Mean | SD[a] |
| S-217622 | >100 | >100 | >100 | >100 | - |
| Remdesivir | >100 | >100 | >100 | >100 | - |

**Table S4.** *In vitro* safety profiles of S-217622 (fumaric acid co-crystal)

| ***In vitro* safety assays** | **Results** |
|---|---|
| hERG inhibition assay | >100 μM |
| Bacterial reverse mutation test (Ames test) | Negative |
| Micronucleus test | Negative |
| 3T3 assay | No Phototoxicity |

**Table S5.** Diffraction data and refinement statistics for SARS-CoV-2 $3CL^{pro}$ complexed with compound **1** and compound **3** (**S-217622**).

a

| | **$3CL^{pro}$ – Compound 1** | **$3CL^{pro}$ – Compound 3 (S-217622)** |
|---|---|---|
| **PDB code** | 7VTH | 7VU6 |
| **Data collection** | | |
| Space group | P21 | P21 |
| Cell dimensions | | |
| *a, b, c* (Å) | 44.41 54.27 114.50 | 55.47 99.23 58.88 |
| *α, β, γ* (°) | 90.00 99.42 90.00 | 90.00 108.05 90.00 |
| Wavelength (Å) | 1.54178 | 1.54178 |
| Resolution (Å) | 28.24 – 2.00(2.05 -2.00) | 28.68 - 1.80(1.84 -1.80) |
| Completeness (%) | 99.5 (98.9) | 99.8(100.0) |
| $R_{merge}$ *(%)*[a,b] | 9.0(27.7) | 6.0(42.9) |
| $I/\sigma(I)$ [a] | 8.3(3.3) | 20.5(2.6) |
| **Refinement** | | |
| Resolution (Å) | 112.952 – 2.001 | 55.986-1.800 |
| No. of reflections | 34366 | 52687 |
| $R_{work}/R_{free}$ | 0.2050/0.2568 | 0.2241/0.2790 |
| *B factor (Å2)* | | |
| Protein | 16.2 | 17.8 |
| Ligand | 27.4 | 14.3 |
| Water | 22.2 | 25.3 |
| R.m.s deviations | | |
| Bond length (Å) | 0.0128 | 0.0175 |
| Bond angles (°) | 1.576 | 1.796 |
| Ramachandran plot (%) | | |
| Favored | 97.2 | 97.5 |
| Allowed | 2.1 | 2.2 |
| Outliers | 0.7 | 0.3 |

Values in parentheses are for the highest resolution shell.
[b] $R_{merge} = \Sigma| I - \langle I \rangle |/\Sigma I$, where *I* is the intensity of observation *I* and $\langle I \rangle$ is the mean intensity of the reflection.
